# Lysine methylation shields an intracellular pathogen from ubiquitylation

**DOI:** 10.1101/2020.11.20.392290

**Authors:** Patrik Engström, Thomas P. Burke, Anthony T. Iavarone, Matthew D. Welch

**Affiliations:** Department of Molecular and Cell Biology, University of California, Berkeley, CA 94720, USA; QB3/Chemistry Mass Spectrometry Facility, University of California, Berkeley, CA 94720, USA

**Author notes:** Correspondence to (P.E.), (M.D.W).

## Abstract

Many intracellular pathogens avoid detection by their host cells. However, it remains unknown how they avoid being tagged by ubiquitin, an initial step leading to anti-microbial autophagy. Here, we show that the intracellular bacterial pathogen *Rickettsia parkeri* uses two protein-lysine methyltransferases (PKMTs) to modify outer membrane proteins (OMPs) and prevent their ubiquitylation. Mutants deficient in the PKMTs were avirulent in mice and failed to grow in macrophages due to ubiquitylation and autophagy. Analysis of the lysine*-*methylome revealed that PKMTs modify a subset of OMPs by methylation at the same sites that are recognized by host ubiquitin. These findings show that lysine methylation is an essential determinant of rickettsial pathogenesis that shields bacterial proteins from ubiquitylation to evade autophagic targeting.

## Main

Intracellular pathogens generally evade the host immune surveillance machinery. This includes avoidance of surface targeting by the host ubiquitylation machinery and subsequent formation of a polyubiquitin coat, a first step in cell-autonomous immunity (*1-5*). The ubiquitin coat recruits autophagy receptors that engage with the autophagy machinery to target cytosol-exposed microbes for destruction (*1, 4-8*). Bacterial outer membrane proteins (OMPs) are targets for the host ubiquitylation machinery (*9, 10*), and the tick-borne obligate intracellular pathogen *Rickettsia parkeri* requires the abundant surface protein OmpB to protect OMPs from ubiquitylation (*11*). However, the detailed mechanisms that *R. parkeri* and other pathogens use to block lysine ubiquitylation of surface proteins, including OMPs, are unknown. We hypothesized that cell surface structures or modifications could provide protection at the molecular level. One such modification is lysine methylation, which is widespread in all domains of life. This modification involves the transfer of one, two, or three methyl groups to the amino group of lysine side chains, the same amino group that can also be modified by ubiquitin (*12*). Whether lysine methylation of bacterial surfaces prevents host detection and promotes intracellular survival has not been explored.

To identify bacteria-derived surface modifications that protect against ubiquitin coating, we screened a library of *R. parkeri* transposon mutants (*13*) (**Table S1**) for increased polyubiquitylation (pUb) relative to wild-type (WT) in Vero cells (an epithelial cell line commonly used to propagate and study intracellular pathogens) by immunofluorescence microscopy (**Fig. 1A**). We subsequently analyzed individual mutants and identified 4 that were ubiquitylated, similar to *ompB* mutant bacteria (*11*) (**Fig. 1B** and **D**). Importantly, these 4 strains expressed OmpB (**Fig. S1**). Two of the strains had insertions in the protein-lysine methyltransferase genes *pkmt1* and *pkmt2*, which are located in two distinct chromosomal regions. The remaining two strains had insertions in the *wecA* and *rmlD* genes, which are required for the biosynthesis of O-antigen (**Fig. S2**), a common surface molecule in Gram-negative bacteria. The O-antigen also protects *Francisella tularensis* from ubiquitylation (*5*), suggesting it performs a conserved function in this process. As a control, we analyzed a strain with a mutation in the *mrdA* gene, which is required for peptidoglycan biosynthesis and cell shape in other bacteria (*14*). This mutant strain had altered shape but was not polyubiquitylated (**Fig. 1B** and **D**), suggesting that not all bacterial cell envelope structures are required to avoid ubiquitylation. In addition, we further quantified the pUb levels and observed that the *pkmt1*::tn mutant, followed by the *ompB* and *pkmt2*::tn mutants, had the highest levels (**Fig. 1E**). These data indicated that OmpB, PKMTs, and the O-antigen protect *R. parkeri* from ubiquitylation.

**Fig 1.**
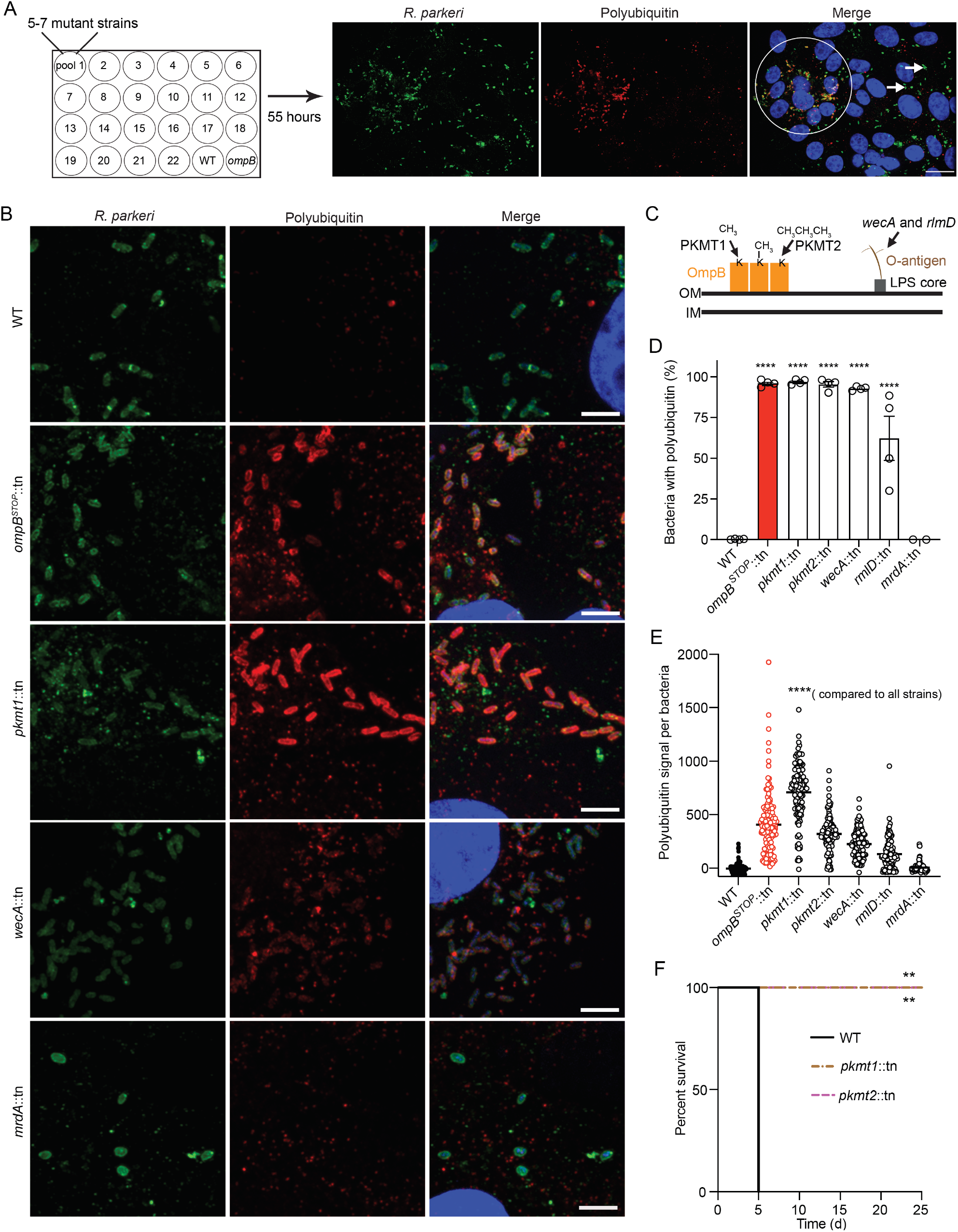
The O-antigen and lysine methylation are virulence factors that protect *R. parkeri* from ubiquitylation. (**A**) Pools of *R. parkeri* (green) mutants screened for increased pUb (red) via immunofluorescence microscopy. DNA, blue. *ompB* mutant (*11*), positive control; WT, negative control. Circle, pUb-positive bacteria; arrows pUb-negative bacteria. Scale bar, 20 μm. (**B**) Infected cells at 72 hours post-infection (h.p.i.) stained as in **A**. Scale bar, 5 μm (*n* = 4). (**C**) Biological function of the genes identified. (**D**) Percentage of bacteria co-localized with pUb at 72 h.p.i. (Data are the mean ± s.e.m.; *n* = 4 for WT and mutant bacteria; *mrdA*::tn; *n* = 2). Statistical comparisons between WT and mutants were performed using a one-way ANOVA with Dunnett’s post-hoc test; **** *P* < 0.0001. (**E**) pUb signal per bacteria. (Lines indicate the means; *n* = 3 fields of vision). Statistical comparisons were performed using a Kruskal-Wallis test with Dunn’s post-hoc test; **** *P* < 0.0001. (**F**) Survival of *Ifnar*^*-/-*^*Ifngr*^*-/-*^ mice intravenously infected with 5×10^6^ WT or mutant bacteria (*n* = 5 mice, WT; *n* = 6 mice, *pkmt1*::tn and *pkmt2*::tn). Statistical comparison between WT and mutants was performed using a two-way ANOVA; ** *P* < 0.01.

The O-antigen was previously shown to be required for rickettsial pathogenesis (*15*), and OmpB was previously found to be required for *R. parkeri* to cause lethal disease in *Ifnar*^-/-^*Ifngr*^-/-^ mice lacking the type I interferon receptor (IFNAR) and IFN-g receptor (IFNGR) (*16*). We therefore examined whether PKMT1 or PKMT2 are important for causing disease *in vivo*. We observed that *Ifnar*^-/-^*Ifngr*^-/-^ mice succumbed to infection with WT but not to *pkmt1*::tn or *pkmt2*::tn bacteria (**Fig. 1F**). Mice infected with the *pkmt1*::tn mutant showed no signs of disease, whereas mice infected with *pkmt2*::tn showed a transient loss in body weight (**Fig. S3**). This indicates that PKMT1 and PKMT2 are determinants of *R. parkeri* pathogenesis.

Because PKMT1 or PKMT2 had previously been shown to methylate OmpB *in vitro* (*17-20*), we next examined whether methylation protects OmpB, or another abundant outer membrane protein OmpA (*21*), from ubiquitylation. First, Vero cells overexpressing 6xHis-tagged ubiquitin were infected with WT and the *ompB, pkmt1*::tn, and *pkmt2*::tn strains. 6xHis-tagged ubiquitin was recruited to the surface of all of the mutants, but not WT bacteria, as observed by immunofluorescence microscopy (**Fig. 2A**). Then, 6xHis-ubiquitylated proteins were affinity-purified from infected cells, and OmpA and OmpB were detected by western blotting. OmpA was shifted towards higher molecular weights in cells infected with mutant, but not WT bacteria, indicating OmpA is ubiquitylated (**Fig. 2B**). Similarly, in comparison with WT bacteria, OmpB was also shifted towards higher as well as lower molecular weights in cells infected with the *pkmt1*::tn and *pkmt2*::tn mutants, suggesting that OmpB is ubiquitylated (**Fig. 2B**). To confirm that methylation protects OmpB and OmpA from ubiquitylation on the bacterial surface, we performed pUb-enrichments of surface fractions from purified bacteria followed by western blotting. This revealed that both OmpB and OmpA shifted towards higher molecular weights in the methyltransferase mutants but not in WT bacteria (**Fig. 2C**). Thus, methylation is critical to protect OMPs from ubiquitylation on the bacterial surface.

**Fig 2.**
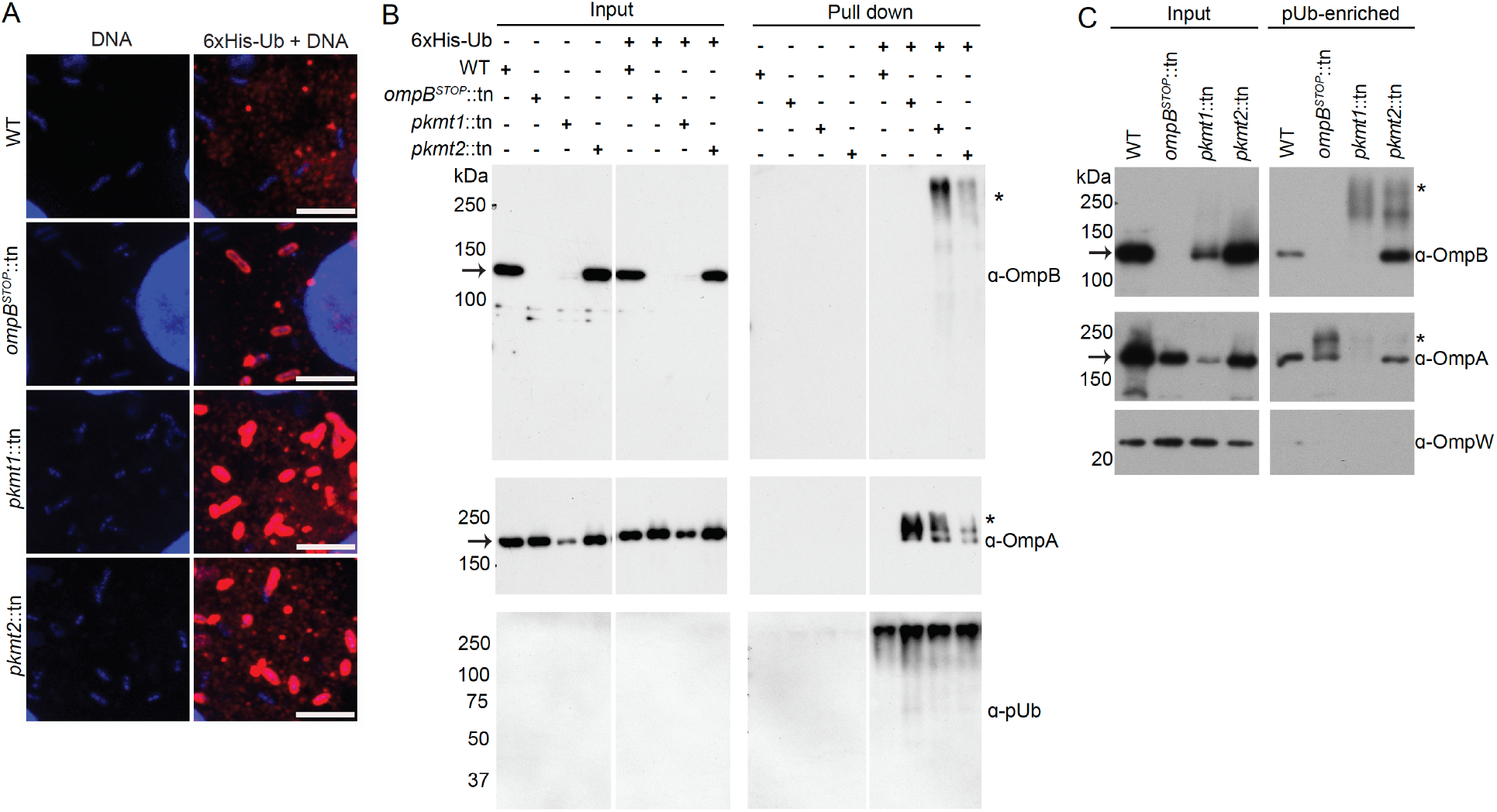
Lysine methylation protects OMPs from ubiquitylation. (**A**) Micrographs of infected Vero cells expressing 6xHis-ubiquitin stained with anti-His antibody (red) and Hoechst (blue, bacterial and host DNA), at 28 h.p.i. Scale bar, 5 μm (representative of *n* = 2). (**B**) Western blot of His-Ub input and pull-down samples from infected control and 6xHis-ubiquitin expressing cells, probed for OmpB, OmpA, and pUb (representative of *n* = 3). (**C**) pUb-enriched (TUBE-1, pan specific) samples from purified bacteria probed for OmpB, OmpA, and OmpW (OmpB and OmpA of endogenous molecular weight represent non-specific binding to TUBE-1 beads) (representative of *n* = 3). Asterisks indicate OmpB and OmpA that exhibit increased molecular weight, indicating ubiquitylation. Arrows indicate OmpB and OmpA of endogenous molecular weight.

Based on our observation that methylation protects both OmpA and OmpB from ubiquitylation, we set out to determine how frequently, and to what extent, lysines of *R. parkeri* OMPs are methylated. Peptides with methylated lysines from whole WT bacteria were quantified using label-free liquid chromatography-mass spectrometry (LC-MS). We then analyzed lysine methylation frequency in abundant OMPs as well as other abundant *R. parkeri* proteins (**Table S2**). This analysis revealed that *R. parkeri* OmpB, OmpA, and surface cell antigen 2 (Sca2) proteins had the highest abundance of methylated peptides. Lysine methylation was also detected in the outer-membrane assembly protein BamA and in a predicted outer membrane protein porin (WP_014410329.1; from here on referred to as OMP-porin) (**Fig. 3A** and **Table S3**). We next mapped both methylated and unmethylated lysines on the above-mentioned OMPs and found that more than 50% of lysines detected from OmpB and OmpA were methylated, and a significant fraction of lysines were also methylated in Sca2 (31%), OMP-porin (30%), and BamA (27%) (**Fig. 3B** and **Fig. S4**). Thus, in *R. parkeri*, lysine methylation of OMPs is common.

**Fig 3.**
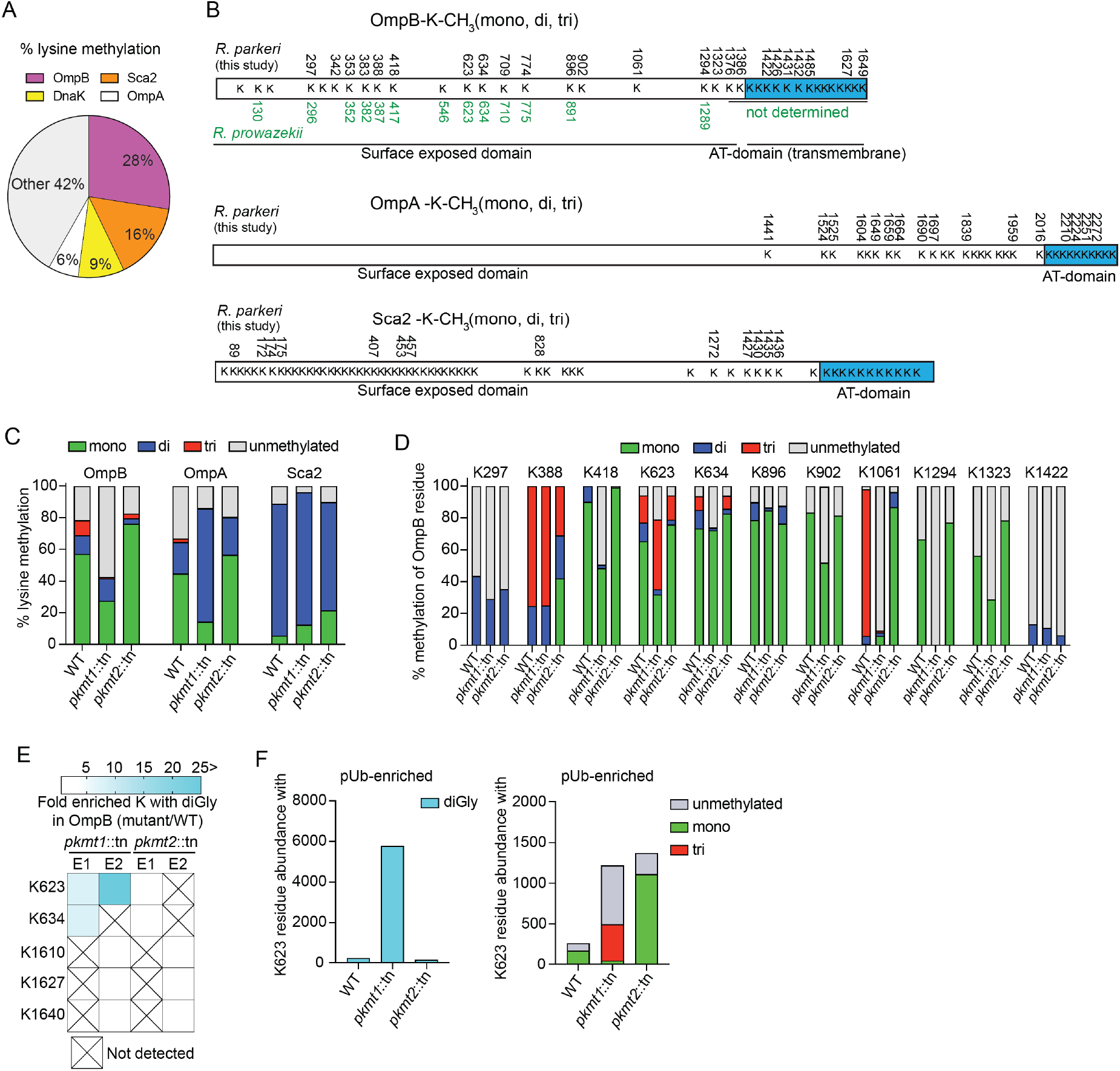
Methylation camouflages lysines from ubiquitylation. (**A**) Percentage lysine methylation abundance of total among the abundantly detected proteins in WT *R. parkeri* determined by LC-MS (data are combined from *n* = 2). (**B**) Methylated lysines (K) are indicated with residue number in each OMP, and lysines in *R. prowazekii* OmpB known to be methylated in green (*17*). K78, K131, K149, K312, and K547 in OmpB were unmethylated (*n* = 5, using data-independent and data-dependent acquisition modes). (**C**) Percentage of total abundances of the lysines methylated in **B**, that is unmethylated, mono-, di-, or tri-methylated (data are the mean of *n* = 2). (**D**) Percentage of individual lysines in OmpB that are unmethylated, mono-, di- or tri-methylated. Only residues repeatedly detected in all strains were analyzed (data are the mean of *n* = 2). (**E**) Heat map representation of OmpB residues from *pkmt1*::tn or *pkmt2*::tn that have similar levels of K-diGly peptides (white boxes) compared to WT, or with a 5-fold, or more, increase of K-diGly peptides (cyan) after pUb-enrichments (*n* = 2). (**F**) Abundances of ubiquitylated (diGly), unmethylated, mono-, di-, or tri-methylated OmpB K623 (data are the mean of *n* = 2). Each experiment (*n*) was performed in technical triplicate.

To identify OMPs that are methylated by PKMT1 and PKMT2 during infection, we compared lysine methylation frequencies in WT with those of *pkmt1*::tn or *pkmt2*::tn mutant bacteria using LC-MS. We found that monomethylation of OmpB, OmpA, the predicted OMP-porin, and another surface cell antigen protein Sca1 were reduced in *pkmt1*::tn compared to WT bacteria (**Fig. 3C** and **Fig. S5**). Dimethylation of rickettsial surface proteins was not reduced in the mutants (**Fig. S6**). Although trimethylation was rare and therefore difficult to analyze at the individual protein level (**Fig. S6**), OmpB had reduced trimethylation levels in both methyltransferase mutants (**Fig. 3C**). Notably, the frequency of unmethylated lysines in OmpB was specifically increased in *pkmt1*::tn bacteria (**Fig. 3C** and **Fig. S5**). Lysine methylation of five other surface proteins (Sca1, Sca2, BamA, LomR, and Pal-lipoprotein), and 21 of 23 abundant proteins with different predicted subcellular distributions, was not affected by mutations in the *pkmt1* or *pkmt2* genes (**Fig. S5**). These data indicate that the PKMTs are required for methylation of a specific subset of OMPs including OmpB.

To determine which OmpB residues are modified by PKMT1 and PKMT2 during *R. parkeri* infection, we analyzed the methylation frequency of individual lysines in the mutants compared to WT using LC-MS. We observed reduced monomethylation frequencies of OmpB K418, K623, K634, K902, K1061, K1294, and K1323 in *pkmt1*::tn bacteria compared with WT, and reduced trimethylation frequencies on K1061 and K388 in the *pkmt2*::tn strain (**Fig. 3D**). These data indicate that several lysines in *R. parkeri* OmpB are methylated by PKMT1 and PKMT2 during infection. Although these results were consistent with previous biochemical results indicating that PKMT1 monomethylates and PKMT2 trimethylates OmpB’s lysines (*17, 19*), we found that total OmpB-methylation is unaffected in the *pkmt2*::tn mutant. This suggests that PKMT1 is a primary methyltransferase for OmpB and that it can compensate, at least partly, for a deficiency in PKMT2.

To test the hypothesis that methylation of specific lysines in OmpB shields the same residues from ubiquitylation, we performed pUb-enrichments of bacterial surface fractions followed by LC-MS to quantify lysines with diglycine (diGly) remnants, a signature for ubiquitin after trypsin digestion. A prediction of this hypothesis is that individual lysines that are heavily methylated in OmpB of *Rickettsia* species (*17*), including WT *R. parkeri* (**Fig. 3D**), are targets for ubiquitylation in *pkmt1*::tn bacteria. In support of this hypothesis, OmpB K634 and K623 in *pkmt1*::tn exhibited 7 to 10,000-fold increased ubiquitylation compared with WT bacteria (**Fig. 3E, F** and **Fig. S7**; **Table S4**). Furthermore, we observed a 13-fold increase in ubiquitylation of the OMP-porin in *pkmt1*::tn bacteria (**Table S4**), indicating that methylation also protects additional OMPs. However, in the *pkmt2*::tn mutant, differential OMP-ubiquitylation was below detection limits (**Fig. 3E, F** and **Table S4**). Together, these data indicate that methylation by PKMT1 camouflages lysines in OMPs from ubiquitylation.

Because polyubiquitylation promotes recruitment of the autophagy receptors p62/SQSTM1 and NDP52 (*7, 8*), we hypothesized that lysine methylation shields OMPs from ubiquitylation to block recruitment of these proteins. Consistent with this hypothesis, we observed that the majority of *pkmt1*::tn, *pkmt2*::tn, as well as *ompB* mutant bacteria, co-localized with p62 and NDP52 by immunofluorescence microscopy (**Fig. S8**). These data demonstrate that PKMTs protect *R. parkeri* from autophagy recognition.

Many pathogenic bacteria including *R. parkeri* grow in immune cells such as macrophages (*11*), despite the fact that microbial detection in such cells triggers anti-bacterial pathways. We therefore investigated whether PKMT1 or PKMT2 were required for evading autophagy targeting and bacterial growth in cultured bone marrow-derived macrophages (BMDMs), as was observed for OmpB (*11*). BMDMs were generated from control mice and mice lacking the gene encoding for autophagy related 5 (ATG5), a protein required for optimal membrane envelopment around pathogens targeted by autophagy, and for their subsequent destruction (*6*). We observed that *pkmt1*::tn mutant bacteria were unable to grow in control BMDMs (*Atg5*^*flox/flox*^), and that growth was rescued in *Atg5*-deficient BMDMs (*Atg5*^-/-^). Further, >91% of *pkmt1*::tn bacteria were labeled with both pUb and p62 in *Atg5*-deficient BMDMs (**Fig. 4A, B, C** and **D**), suggesting that ubiquitin-tagged bacteria were not restricted when the autophagy cascade was prevented. In contrast, the *pkmt2*::tn mutant did not have a major growth defect compared with WT bacteria and 50% of the bacteria were labeled with p62, irrespective of host genotype (**Fig. 4A, B, C** and **D**), consistent with less pronounced ubiquitylation phenotypes compared to *pkmt1*::tn bacteria. Collectively, these data indicate that methylation is required for *R. parkeri* growth in macrophages by avoiding autophagic targeting.

**Fig 4.**
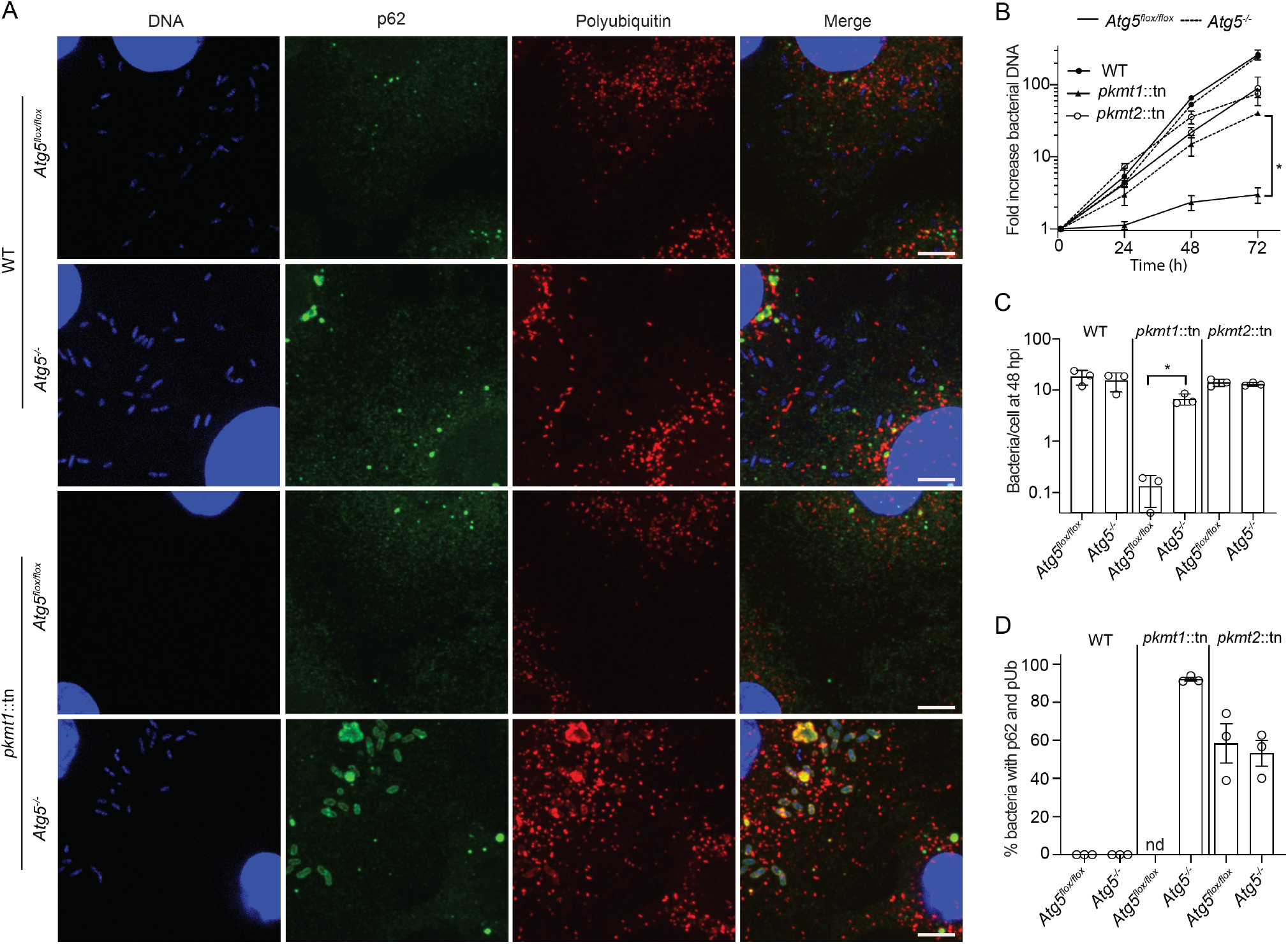
Methylation prevents ATG5-dependent *R. parkeri* killing in macrophages. (**A**) Micrographs of infected control (*Atg5*^*flox/flox*^) and *Atg5*-deficient (*Atg5*^-/-^) BMDMs at 48 h.p.i., stained for DNA (blue), pUb (red), and p62 (green). Arrow indicate a bacterium positive for both pUb and p62. Scale bars, 5 μm (*n* = 3). (**B**) Bacterial growth curves in control and *Atg5*^-/-^ BMDMs, as measured by genomic equivalents using qPCR (*n* = 3). (**C**) Quantification of the mean number of bacteria per cell (*n* = 3). (**D**) Quantification of the percentage of bacteria with p62 and pUb (*n* = 3). nd, not determined. Statistical comparisons in **B** and **C** were performed using a Brown-Forsyth and Welch ANOVA with Dunnett’s post-hoc test; *, *P* < 0.05. Presented data are the mean ± s.e.m.

Our work reveals a detailed molecular mechanism that camouflages bacterial surface proteins from host detection. In particular, we found that lysine methylation was essential for blocking ubiquitylation, a first step in cell-autonomous immunity (*1-4*). This highlights an intricate evolutionary arms race between pathogens and hosts and reveals a strategy that pathogens can adapt to counteract host responses. The lysine methyltransferases PKMT1 and PKMT2 are conserved between rickettsial species (**Fig. S9** and **Fig. S10**) and contain a core Rossmann fold found in the broader superfamily of class I methyltransferases that exist in diverse organisms (*12, 18*). Thus, we propose that lysine methylation, and potentially other lysine modifications, could be used by pathogens, symbionts, and perhaps even in eukaryotic organelles, to prevent undesirable surface ubiquitylation and downstream consequences including elimination by autophagy. Further study of microbial surface modifications will continue to enhance our understanding of the pathogen-host interface and could ultimately lead to new therapeutic targets for treating human diseases including those caused by infectious agents.

## Acknowledgements

We thank Prof. Dan Portnoy (UC Berkeley) and Dr. Lilliana Radoshevich (University of Iowa) for providing comments on this manuscript, and Neil Fischer (UC Berkeley) for critically reading the manuscript. We also thank Dr. Hwan Kim (Stony Brook University) for kindly providing the O-antigen antibody and for fruitful discussions. We are also grateful for the *Atg5*^*flox/flox*^ and *Atg5*^*-/-*^ BMDMs provided by Dr. G. Golovkine (UC Berkeley), and Profs. Jeffery S. Cox and Michael Rape (both UC Berkeley) for fruitful discussions.

## Author contributions

P.E. conceived the study with the assistance of M.D.W. P.E. performed laboratory work and analysis, except for mass spectrometry, which was conducted together with A.T.I, and animal experiments, which were conducted by T.P.B. P.E. drafted the initial manuscript, and A.T.I, T.P.B, and M.W.D provided editorial feedback.

## Data and materials availability

All data used in the analysis are provided in the main text or supplementary materials. Materials are available upon request.

## Supplementary Figures

**Fig S1.**
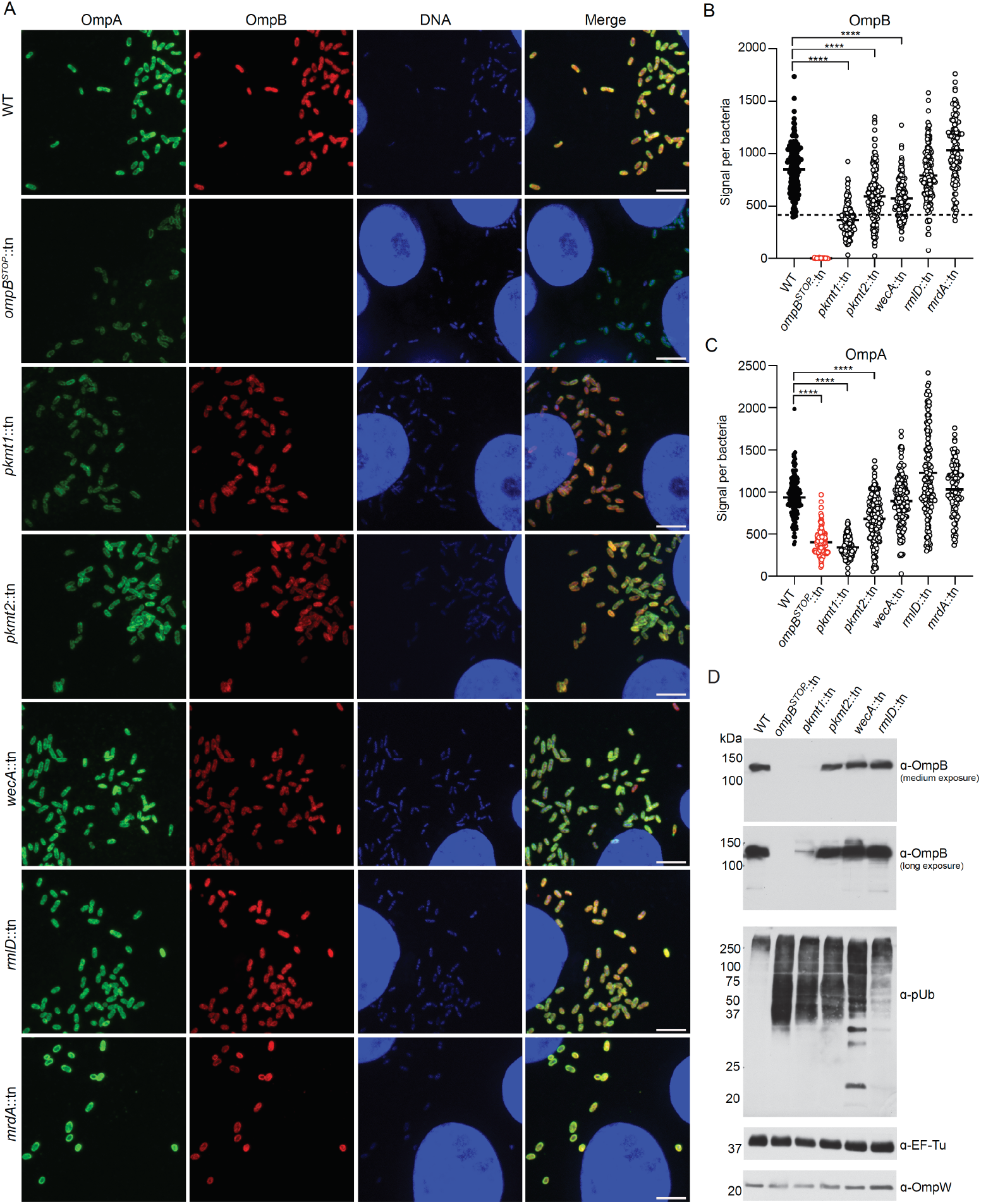
Polyubiquitylated mutant *R. parkeri* strains are positive for OmpA and OmpB. **(A**) Micrographs of Vero cells infected with the indicated strains stained for OmpA (green, 13-3 antibody), OmpB (red, OmpB-antibody), and DNA (blue, Hoechst) at 72 h.p.i. (representative of *n* = 3). Scale bar 5 μm. (**B**) Quantification of OmpB signal per bacteria. (Lines indicate the means; *n* = 3 fields of vision; ≥ 50 bacteria per field of vision were analyzed). Statistical comparisons were performed using a Kruskal-Wallis test with Dunn’s post-hoc test; **** *P* < 0.0001 between indicated strains. Dashed line indicates that the majority of mutant bacterial populations, except *pkmt1*::tn bacteria, have OmpB levels comparable to WT bacteria. (**C**) Quantification of OmpA signal per bacteria as in **B**. (**D**) Western blot of 5×10^6^ purified bacteria probed for OmpB, pUb, OmpW and EF-Tu (bacterial loading controls) (representative of *n* = 2).

**Fig S2.**
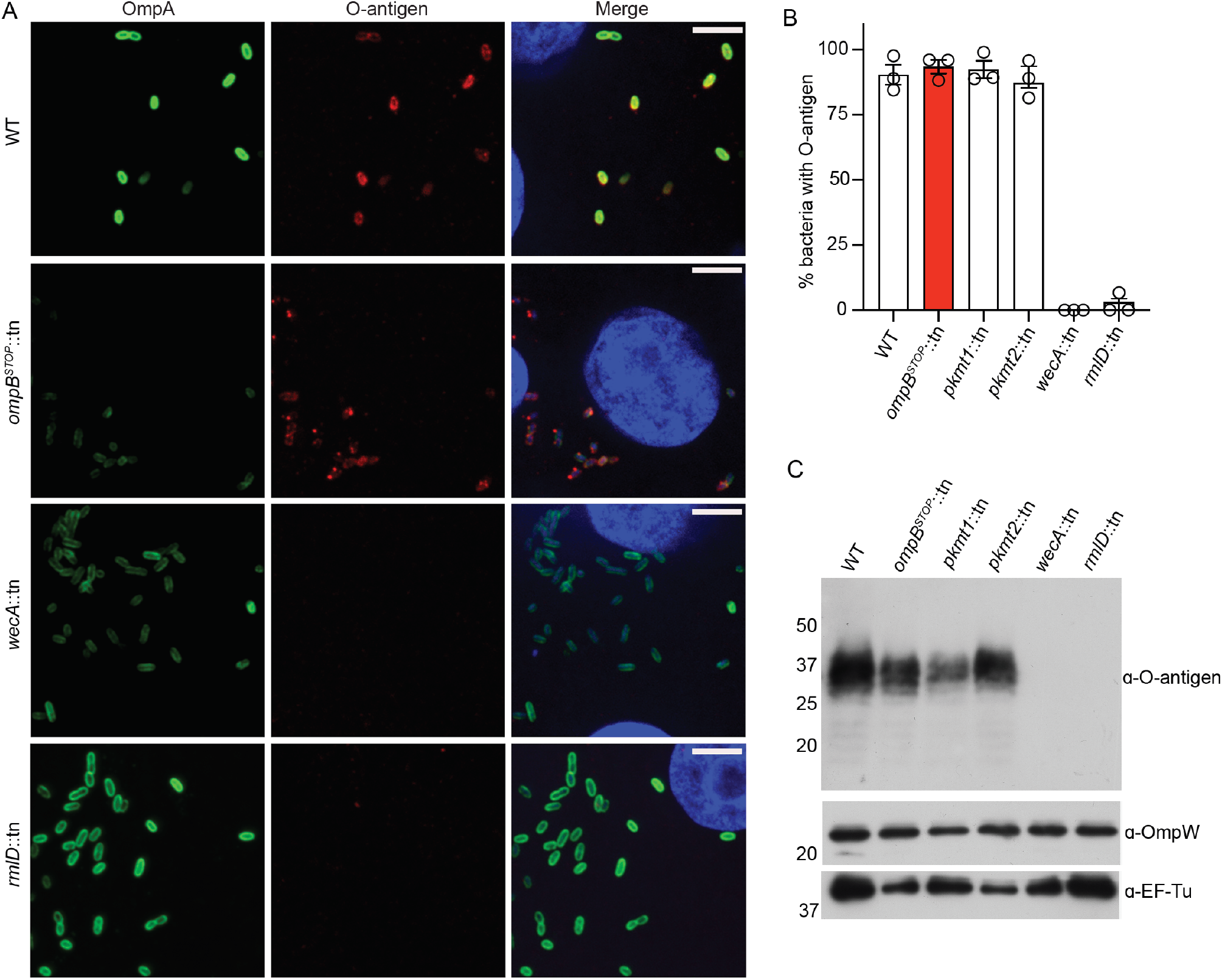
The *wecA*::tn and *rmlD*::tn bacteria lack the O-antigen. (**A**) Micrographs of Vero cells infected with the indicated strains stained for OmpA (green, 13-3 antibody), the O-antigen (red, O-antigen antibody(*15*)) and DNA (blue, Hoechst) at 72 h.p.i. (representative of *n* = 3). Scale bars, 5 μm. (**B**) Percentage of bacteria positive for the O-antigen at 72 h.p.i. *n* = 2; ≥ 90 bacteria were counted in each infection (data are the mean ± s.e.m.; *n* = 3). (**C**) Western blot of 5×10^6^ purified WT and mutant bacteria probed for the O-antigen. OmpW and EF-Tu were used as bacterial loading controls (representative of *n* = 3).

**Fig S3.**
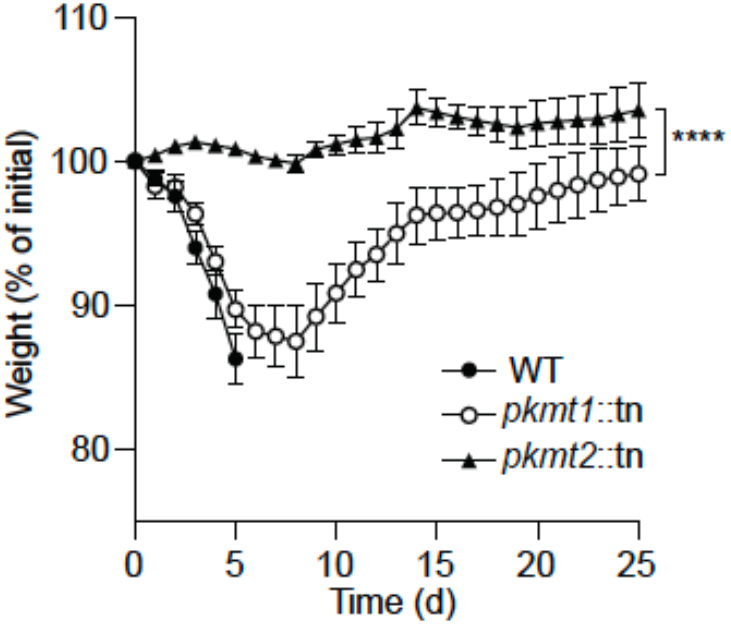
PKMT1 plays a more significant role in causing disease *in vivo* compared to PKMT2. (**A**) Weight changes of *Ifnar*^*-/-*^*Ifngr*^*-/-*^ mice intravenous infected with 5×10^6^ WT, *pkmt1*::tn, or *pkmt2*::tn bacteria (data are the mean ± s.e.m. *n* = 5, WT; *n* = 6, *pkmt1*::tn and *pkmt2*::tn, combined from two independent experiments). A two-way ANOVA from 0 to 25 days post-infection (d.p.i.) was used to statistically compare the weight changes between the *pkmt1*::tn and *pkmt2*::tn mutants. **** *P* < 0.0001.

**Fig S4.**
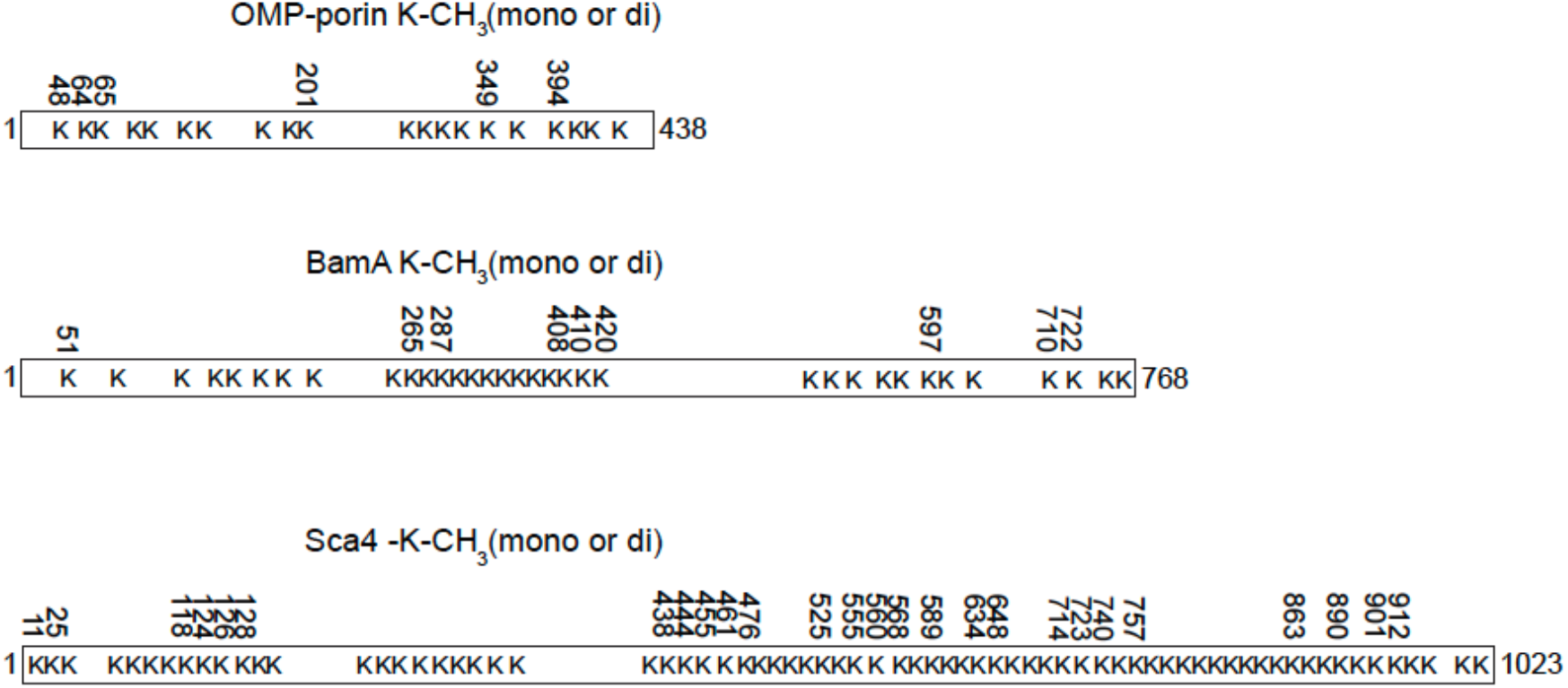
Lysine-methylome reveals that *R. parkeri* OMP-porin, BamA, and the released factor Sca4 are methylated. Methylated lysines are indicated with residue number, and unmethylated residues without, determined by LC-MS as in **Fig. 3B** (data is combined from five independent experiments). See also **Table S2**.

**Fig S5.**
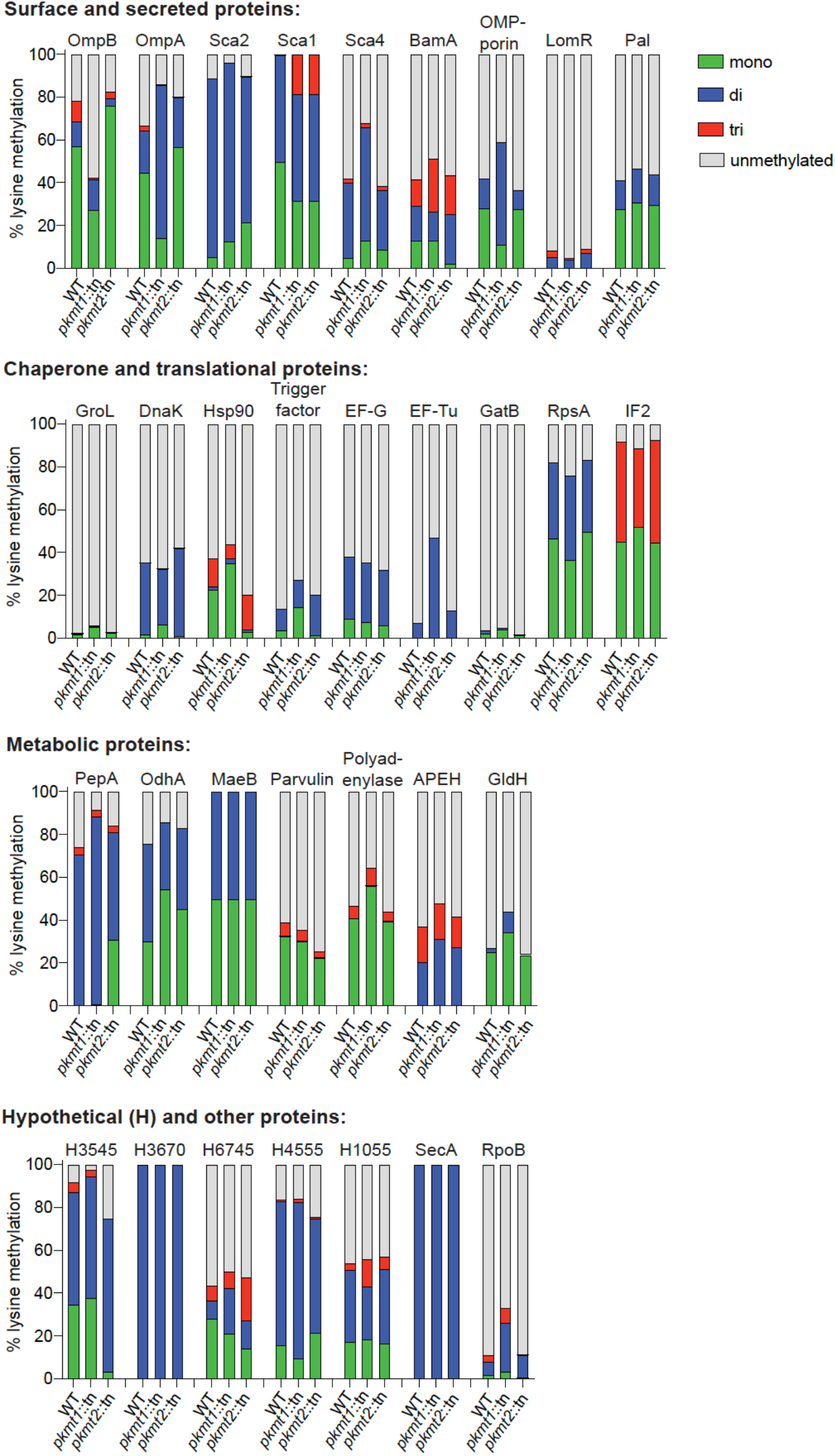
PKMT1 primarily modifies bacterial OMPs. Percentage of total abundances of the lysines methylated in WT, that is unmethylated, mono-, di-, or tri-methylated in respective strain. Proteins with ≥3 lysines methylated detected in independent experiments are shown (mean of *n* = 2, performed in technical triplicates). OmpB, OmpA and Sca2 data are the same as in **Fig. 3C**. See also **Table S2**.

**Fig S6.**
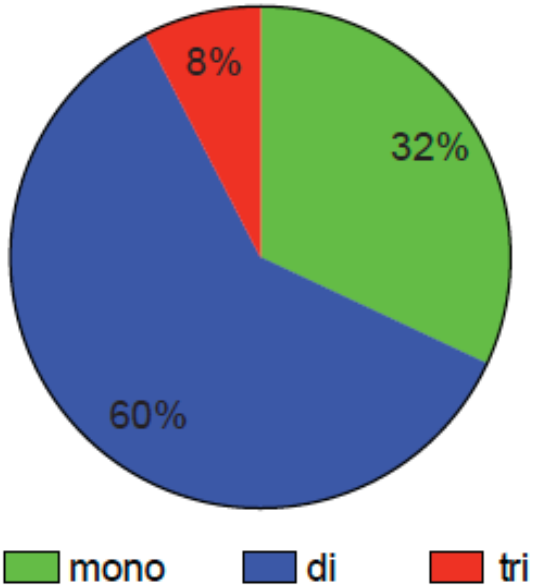
Dimethylation is a common methylation state among the abundant *R. parkeri* proteins. Percentage of total lysines abundances detected to as mono-, di-, or tri-methylated in WT bacteria of the proteins indicated in **Table S2** (mean of *n* = 2, performed in technical triplicates).

**Fig S7.**
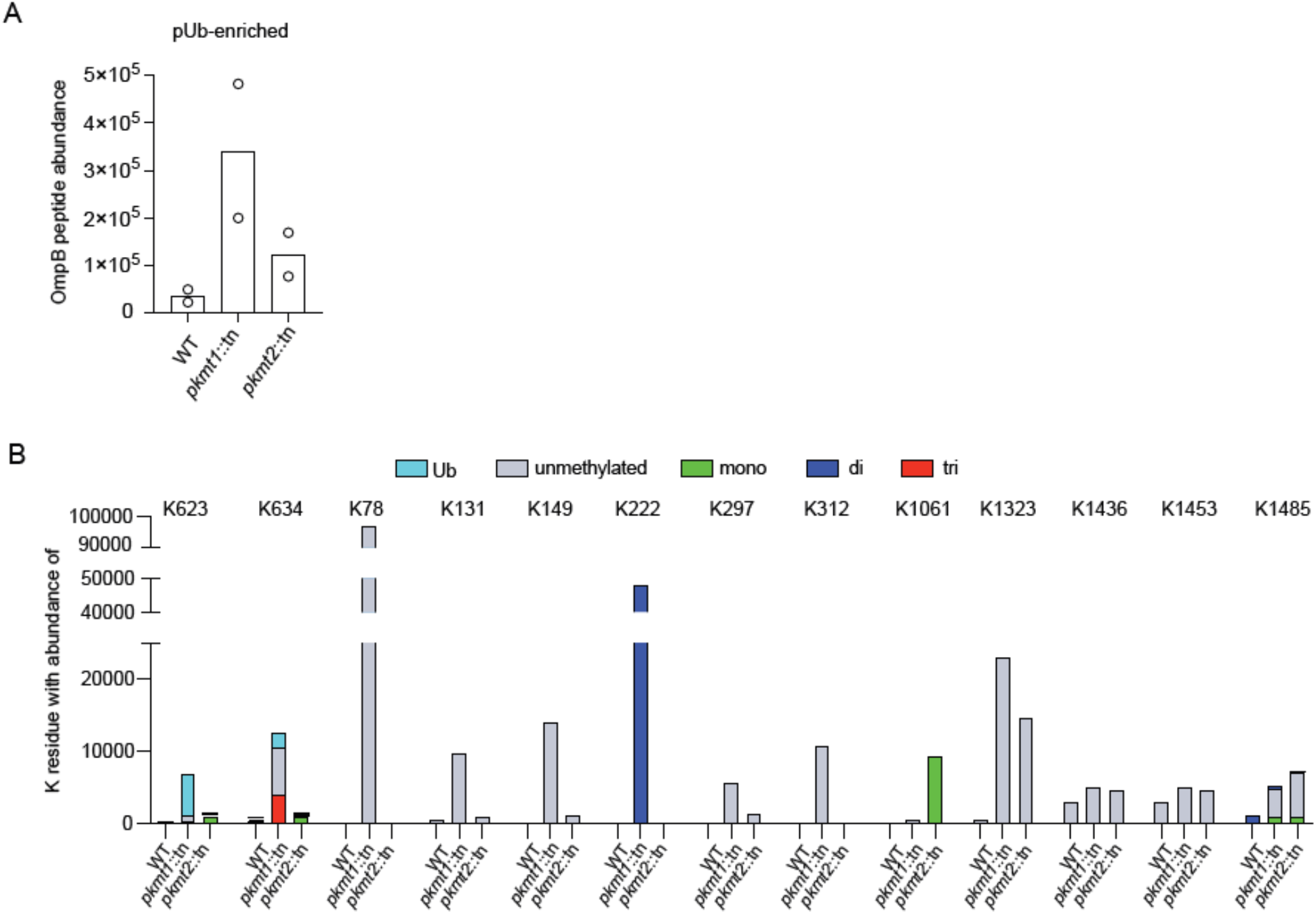
OmpB K623 and K634 are frequently ubiquitylated when PKMT1-mediated methylation is reduced. (**A**) OmpB peptide abundance values after pUb-enrichments from respective strain, as determined by LC-MS (data are mean of *n* = 2, performed in triplicates). (**B**) Abundance values of lysines in OmpB that are ubiquitylated (diGly), unmethylated, mono-, di-, or tri-methylated, in respective strain after pUb-enrichments as in **Fig 2C**. Only residues detected in independent experiments are shown (data are the mean of *n* = 2, performed in technical triplicates). K623 data in **B** are the same as in **Fig. 3F**.

**Fig S8.**
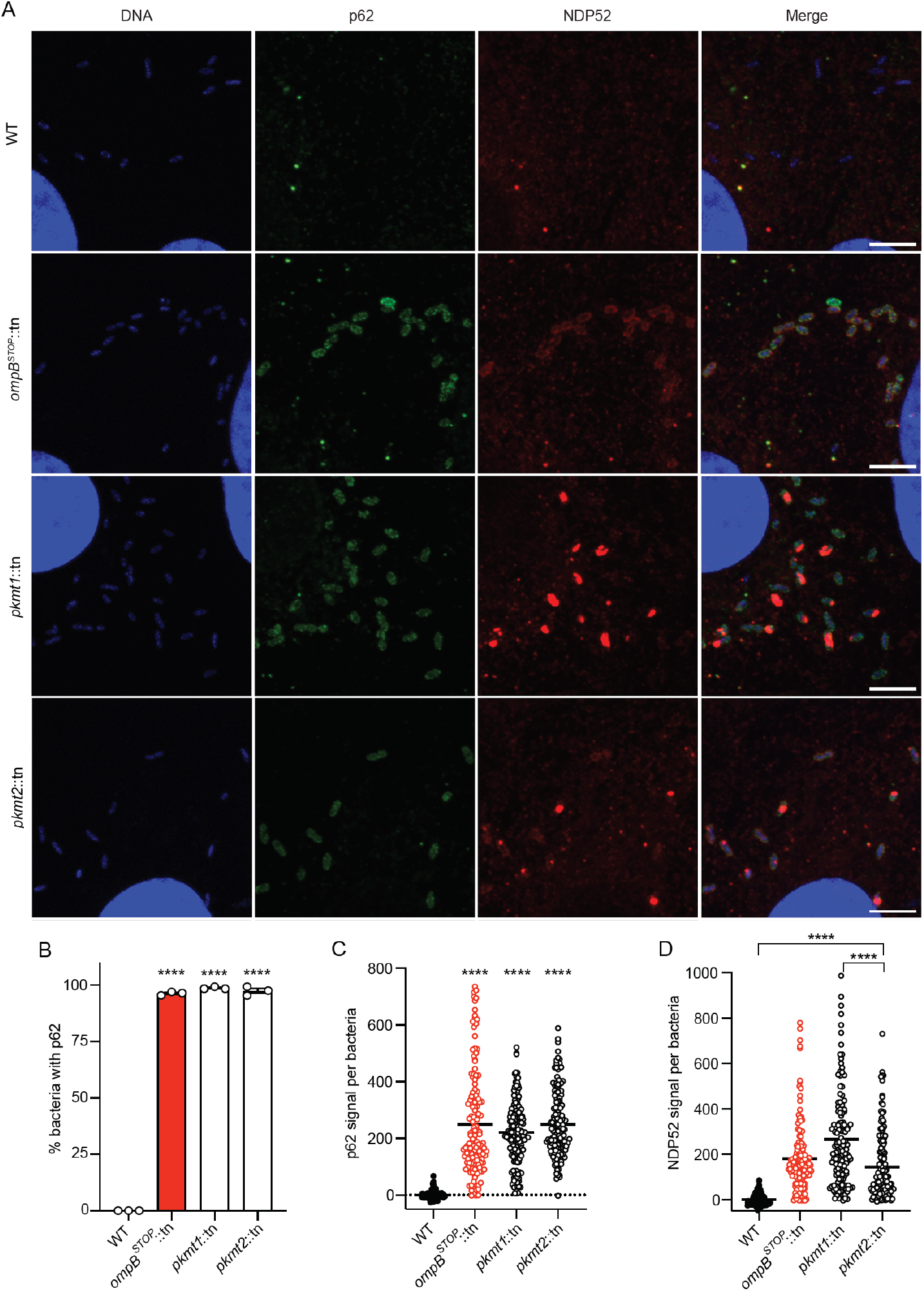
OmpB and lysine methylation block recruitment of autophagy receptors to *R. parkeri*. (**A**) Micrographs of Vero cells infected with the indicated strains: stained for bacterial and host DNA (blue, Hoechst), p62 (green, p62-antibody), and NDP52 (red, NDP52-antibody) at 72 h.p.i (*n* = 4 for p62 staining; *n* = 2 for NDP52 staining). (**B**) Percentage of bacteria that show rim-like surface localization of p62 at 72 h.p.i. (Data are the mean ± s.e.m.; *n* = 3; ≥ 142 bacteria were counted for each infection). Statistical comparisons between WT and *ompB*^*STOP*^::tn, *pkmt1*::tn, *pkmt2*::tn were performed using a one-way ANOVA with Tukey’s post-hoc test; **** *P* < 0.0001. (**C**) Quantification of p62 signal per bacteria from a representative experiment. Lines indicate the means. (*n* = 3 fields of vision; ≥ 50 bacteria per field of vision were analyzed). Statistical comparisons were performed using a Kruskal-Wallis test with Dunn’s post-hoc test; **** *P* < 0.0001 between indicated strains. (**D**) Quantification of NDP52 signal per bacteria as in **C**.

**Fig S9.**
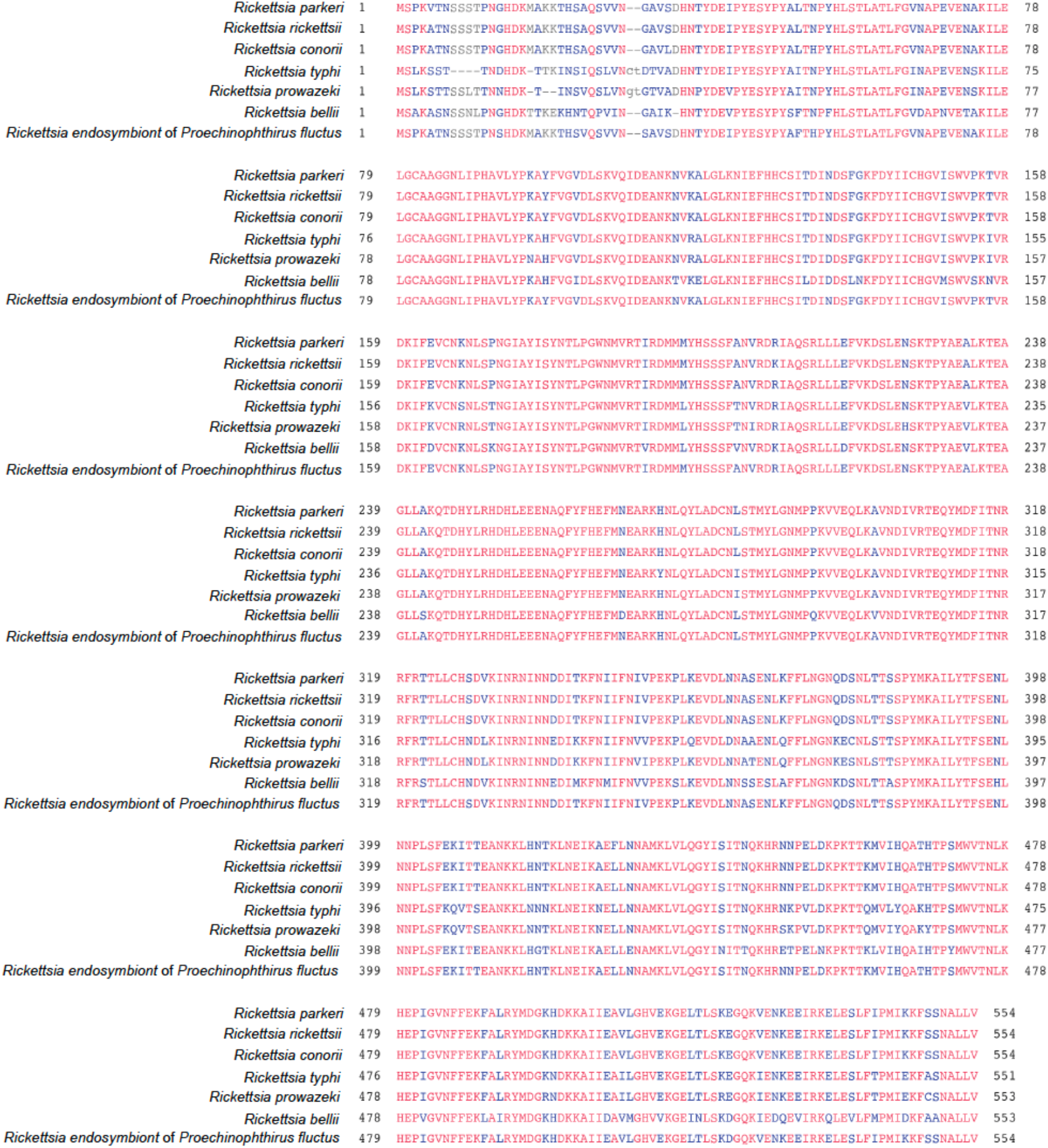
The PKMT1 enzyme is highly conserved between diverse rickettsial species. Amino acid sequence alignment of PKMT1 from *R. parkeri* (<u>WP_014411082.1</u>), *R. rickettsii* (<u>WP_012262600.1</u>), *R. conorii* (<u>WP_016926592.1</u>), *R. typhi* (<u>WP_011191207.1</u>), *R. prowazekii* (<u>WP_004596928.1</u>), *R. bellii* (<u>WP_045799810.1</u>), and a *Rickettsia* endosymbiont (<u>WP_062811822.1</u>), using <u>COBALT</u>. Amino acids indicated in red are identical; blue, variation between species; grey, one or more of the analyzed proteins are lacking this residue(s).

**Fig S10.**
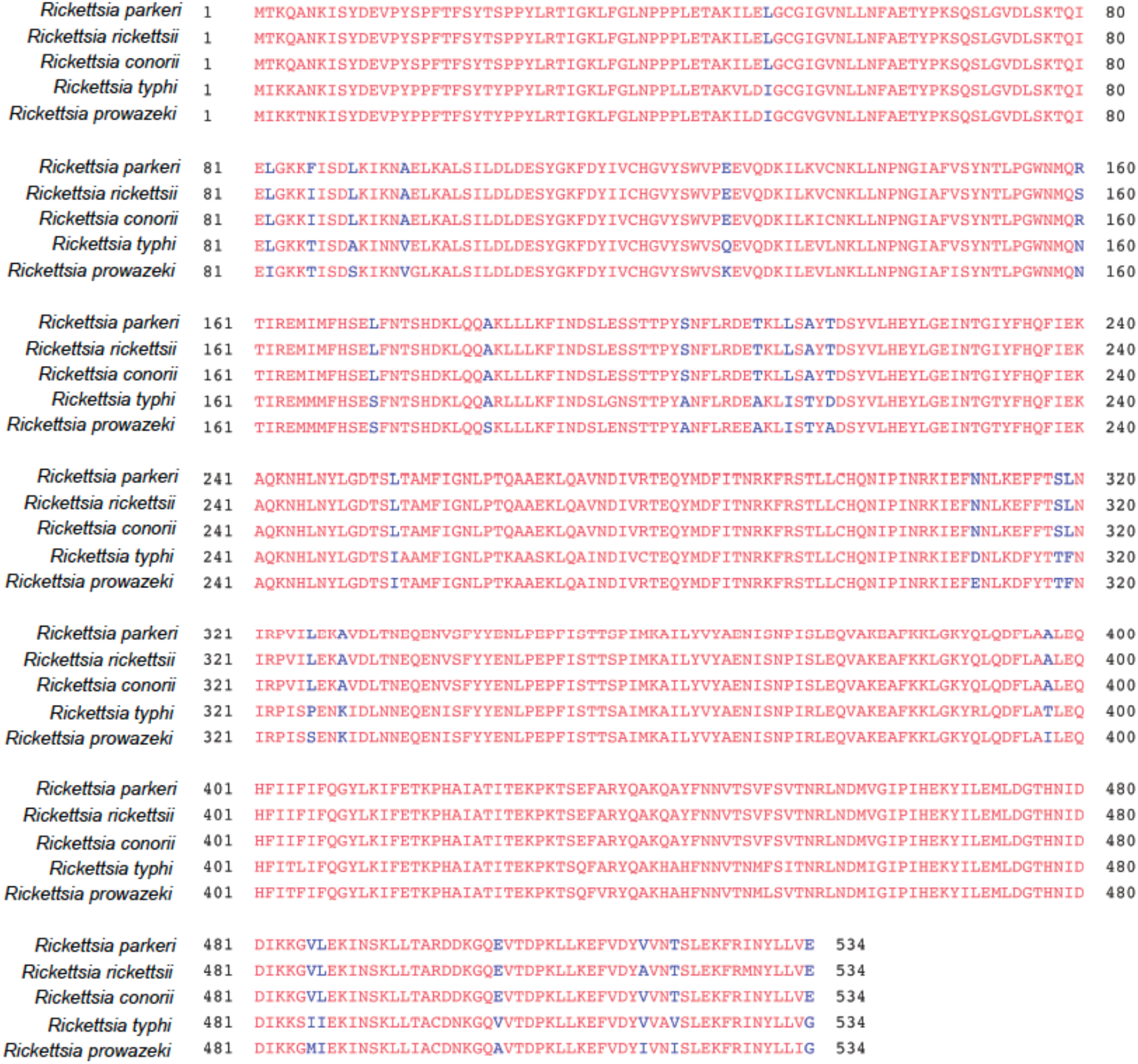
The PKMT2 enzyme is highly conserved between virulent rickettsial species but absent from *R. bellii* and the *Rickettsia endosymbiont Proechinophthirus fluctus*. Amino acid sequence alignment of PKMT2 from *R. parkeri* (<u>WP_014410272.1</u>), *R. rickettsii* (<u>WP_012150259.1</u>), *R. conorii* (<u>WP_016925880.1</u>), *R. typhi* (<u>WP_011190574.1</u>), and *R. prowazekii* (<u>WP_004596662.1</u>) using <u>COBALT</u>. PKMT1 of *R. bellii* and the *Rickettsia endosymbiont Proechinophthirus fluctus* showed 51% identity to PKMT2 of *R. parkeri* using single alignment BLAST at NCBI; however, no PKMT2 variants could be found in these organisms, and therefore they were excluded from this analysis. Amino acids indicated in red are identical; blue, variation between species.

**Fig S11.**
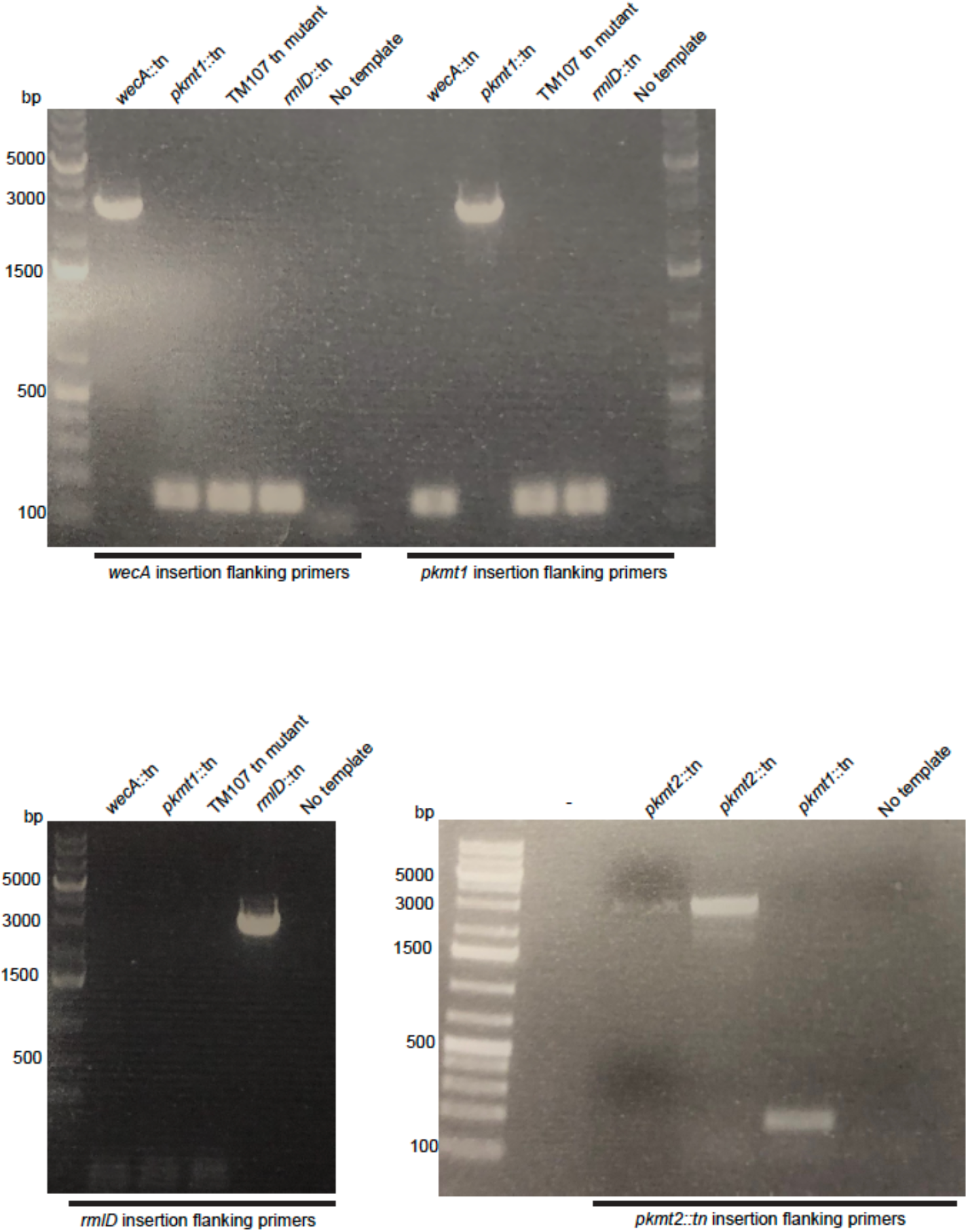
Strain validation of clonality and insertion site using primers for flanking chromosomal regions (*n* = 1).

## Materials and Methods

### Cell lines and primary mouse macrophages

Vero cells were purchased from the UC Berkeley Cell Culture Facility and the identity was repeatedly confirmed by mass-spectrometry analysis. Cells were grown at 37 °C and 5% CO_2_ in DMEM (Gibco, cat. no. 11965) with high glucose (4.5 g l^-1^) and 2% heat-inactivated (30 min, 56 °C, in a water-bath) fetal bovine serum (Gemcell). Vero cells were confirmed to be mycoplasma negative by DAPI staining and fluorescence microscopy screening at the UC Berkeley Cell Culture Facility.

BMDMs generated from the femurs of mutant *Atg5*^*flox/flox*^ and matched *Atg5*^-*/-*^ C57BL/6 mice were a kind gift from the laboratory of Jeffery S. Cox (UC Berkeley), and they were prepared as previously described (*11*) although in the absence of antibiotics. Genotypes were confirmed by PCR and Sanger sequencing at the UC Berkeley DNA Sequencing Facility, as previously described (*11*).

### *Rickettsia parkeri* strain generation and validation

WT *R. parkeri* strain Portsmouth (NCBI accession no. <u>NC_017044.1</u>; originally a gift from C. Paddock, Center for Disease Control and Prevention) and bacterial stocks of *ompB*^*STOP*^::tn (the genome sequences of these bacterial strains are available at the Sequence Read Archive as accession no. <u>SRP154218</u> (WT, <u>SRX4401164</u>; *ompB*^*STOP*^::tn, <u>SRX4401167)</u>), *pkmt1*::tn, *pkmt2*::tn, *wecA*::tn and *rlmD*::tn mutants were propagated and purified every ∼6-10 months as described below. Side-by-side experimental comparisons were made between stocks prepared at similar times.

*R. parkeri pkmt1*::tn, *pkmt2*::tn, *wecA*::tn, and 114 other transposon insertion mutant strains, screened for pUb, were previously isolated in a screen for small plaque mutants (*13, 22*). The *ompB*^*STOP*^::tn was previously isolated in a suppressor screen and lacked expression of OmpB (*11*). The *rmlD*::tn, and 132 other transposon insertion mutant strains, screened for pUb, were isolated in an independent screen in which mutants were isolated without regard for plaque size (**Table S1**). The genomic locations of transposon insertion sites for all mutants were determined by semi-random nested PCR. To verify the insertions and clonality, we used PCR reactions that amplified the transposon insertion site using primers for flanking chromosomal regions: 5’GCTCACTAGATAGCACTCG’3 and 5’GCTCGATTTATCTCACTTTATG’3 for *rlmD*::tn, 5’CGTTTAATAGTCCAGTTAATTTGT’3 and 5’CCGTCTATACCGTCCATAAAAT’3 for *wecA*::tn, 5’GCATCGAAATAACCCTGAG’3 and 5’GCAAACTTCTCAAAGAAATTAACG’3 for *pkmt1*::tn, 5’GCTAAGAAATCTTCTAATTTGATATTTTAC’3 and 5’CGAAAATTTACCTGAGCCTT’3 for *pkmt2*::tn, 5’CGACACATAATAGCACAAACTAC’3 and 5’GCGGAGGCGGTAGTAAAG’3 for *mrdA*::tn (**Fig. S11**).

### Screening for pUb-positive strains

To prepare the mutant library for screening, passage 1 (P1) transposon insertion mutants were amplified one time in Vero cells using 24-well cell culture plates. At 5-12 d.p.i, when 50-70% of the infected cells appeared to be rounded up (as a sign of infection) by visual inspection using a light microscope, cell culture media were completely removed, and cells were subsequently lysed in 500 μL cold sterile water for 2-3 minutes (min). Next, 500 μL of 2x cold sterile brain-heart-infusion (BHI) broth (BD Difco, 237500) was added to the lysed cells, resuspended, and P2 bacteria were transferred to cryogenic storage vials and frozen at −80 °C.

To screen for pUb-positive strains, 10-40 μL of each of five to seven P2 mutant bacterial strains were diluted in 1 mL of room temperature (RT) cell culture media supplemented with 2% FBS. Subsequently, the pooled bacterial suspension was centrifuged at 250*g* for 4 min at RT onto confluent Vero cells grown on coverslips in 24-well plates. Cells were then incubated at 33 °C and fixed at 50-55 h.p.i. with pre-warmed 4% paraformaldehyde (PFA) (Ted Pella Inc., 18505) for 10 min at RT. If cells were over-infected (i.e., individual infection foci had grown together) as determined by immunofluorescence microscopy, infections of that specific pool were repeated using reduced volumes of P2 bacteria. Next, fixed cells were permeabilized with 0.2% Triton-X for 5 min and then stained with the anti-*Rickettsia* I7205 antibody (1:500 dilution; gift from Ted Hackstadt) and anti-polyubiquitin FK1 antibody (Enzo Life Sciences, BML-PW8805-0500; 1:250 dilution), followed by Alexa 488 anti-rabbit antibody (Invitrogen, A11008; 1:500 dilution) or goat anti-mouse Alexa-568 (Invitrogen, A11004). Whole coverslips were manually inspected on a Nikon Ti Eclipse microscope with 60x (1.4 numerical aperture) Plan Apo objective. The initial screen revealed that five out of 39 mutant pools contained pUb-positive areas.

In a secondary screen, individual strains from the pUb-positive pools were used to infect Vero cells, as stated above. Infected cells were also fixed and stained as above except that a post-fixation step using 100% methanol for 5 min was included and an OmpB antibody (*11*) and Hoechst (Sigma, B2261, 1:2500 dilution) was used instead of the anti-*Rickettsia* I7205 antibody. Samples were inspected as above and strains were scored as follows: **1**) pUb-negative (49 strains), **2**) a few infection foci were pUb-positive (2 strains: Sp mutant 24, insertion at bp position 753916; Sp mutant 94, insertion at bp position 774831), **3**) bacteria in the center of foci were pUb-positive but not on the edges (2 strains: Sp mutant 43, insertion at bp position 651602-651604; Sp mutant 45, insertion at bp position 751156), **4**) almost all bacteria in all foci were pUb-positive (4 strains: *pkmt1*::tn, insertion at bp position 1161553 (gene *MC1_RS06185*); *pkmt2*::tn, insertion at bp position 34100 (*MC1_RS00180*); *wecA*::tn, insertion at bp position 1223170 (*MC1_RS06510*); and *rmlD*::tn, insertion at bp position 455753 (*MC1_RS02345*).

### *Rickettsia* purification

*R. parkeri* strains were propagated as described previously (*11*). “Purified bacteria” were from five T175 flasks of Vero cells that after 5-8 days of infection (normally ∼75% infected as observed by light microscopy) were harvested in the media using a cell scraper. Bacteria were tehn centrifuged 12000*g* for 15 min at 4 °C in pre-chilled tubes. Pellets were resuspended in cold K-36 buffer (0.05 M KH_2_PO_4_, 0.05 M K_2_HPO_4_, pH 7, 100 mM KCl and 15 mM NaCl) and a pre-chilled dounce-homogenizer (tight fit) were used for 60 strokes to release bacteria from host cells. The homogenate was then centrifuged at 200*g* for 5 min at 4 °C to remove cellular debris. The supernatant was overlaid onto cold 30% v/v MD-76R (Mallinckrodt Inc., 1317-07) diluted in K-36, and centrifuged at 58300*g* for 20 min at 4 °C in an SW-28 swinging-bucket rotor. The bacterial pellet was resuspended in cold 1x BHI broth (0.5 mL BHI per infected T175 flask), and after letting DNA precipitates sediment to the bottom of the tubes, bacterial suspensions were collected, aliquoted and frozen at −80 °C.

“Gradient-purified bacteria” were from ten T175 flasks of Vero cells, purified as above with the addition of a 40/44/54% v/v MD-76R (diluted in K-36 buffer) gradient step centrifuged at 58300*g* for 25 min at 4 °C using the SW-28 swinging bucket rotor. The bacteria were then collected from the 44-54% interface, diluted in K-36 buffer, and pelleted by centrifugation at 12000*g* for 15 min at 4 °C. The pellet was resuspended in cold 1x BHI broth and subsequently aliquoted and frozen at −80 °C.

### OmpW and EF-Tu antibody production

The sequence encoding amino acids 22-224 of outer membrane protein W (OmpW; WP_014411122.1) (a protein that lacks the signal peptide), or full-length Elongation factor Tu (EF-Tu; WP_004997779.1), were amplified by PCR from *R. parkeri* genomic DNA, and subsequently cloned into plasmid pETM1, which encodes N-terminal 6xHis and maltose-binding proteins (MBP) tags. From the resulting plasmids, fusion proteins were expressed in *E. coli* strain BL21 codon plus RIL-Cam^r^ (DE3) (QB3 Macrolab, UC Berkeley) by induction with 1 mM isopropyl-β-D-thio-galactoside (IPTG) for 2-2.5 hours at 37 °C. Bacterial pellets were resuspended in lysis buffer (50 mM NaH_2_PO_4_, pH 8.0, 300 mM NaCl, 1mM EDTA, and 1 mM dithiothreitol (DTT)) and stored at −80 °C. For protein purification, bacteria were thawed, lysozyme was added to 1 mg/mL (Sigma, L4919), and lysis was carried out by sonication. Lysates containing 6xHis-MBP-OmpW and 6xHis-MBP-EF-Tu were incubated on amylose resin (New England Biolabs, E8021L) (Qiagen, 1018244) and bound proteins were eluted in lysis buffer lacking EDTA and DTT but containing 10 mM maltose. Fractions were analyzed by SDS-PAGE and those with the highest concentrations of fusion proteins were pooled to generate rabbit antibodies against OmpW and EF-Tu. 1.2 mg of purified 6xHis-MBP-OmpW and 6xHis-MBP-EF-Tu proteins were sent to Pocono Rabbit Farm and Laboratory (Canadensis, PA), and immunization was carried out according to their 91-d protocol.

### Western blotting

To determine the levels of bacterial and host proteins in purified bacterial samples, 30%-purified bacterial samples were boiled in 1x SDS loading buffer (150 mM Tris pH 6.8, 6% SDS, 0.3% bromophenol blue, 30% glycerol, 15% β-mercaptoethanol) for 10 min, then 5×10^6^ PFUs were resolved on an 8-12% SDS-PAGE gel and transferred to an Immobilon-FL polyvinylidene difluoride membrane (Millipore, IPEL00010). Membranes were probed for 30 min at room temperature or 4°C overnight with antibodies as follows: affinity-purified rabbit anti-OmpB antibody (*11*) diluted 1:200-30000 in TBS-T (20 mM Tris, 150 mM NaCl, pH 8.0, 0.05% Tween 20 (Sigma, P9416)) plus 5% dry milk (Apex, 20-241); mouse monoclonal anti-OmpA 13-3 antibody diluted 1:10000-50000 in TBS-T plus 5% dry milk; rabbit anti-OmpW serum diluted 1:8000 in TBS-T plus 5% dry milk; mouse monoclonal FK1 anti-polyubiquitin antibody diluted 1:2500 in TBS-T plus 2% BSA; rabbit anti-EF-Tu serum diluted 1:15000 in TBS-T plus 5% dry milk; or rabbit anti-O-antigen serum 1:5000 in TBS-T plus 5% dry milk. Secondary antibodies were: mouse anti-rabbit horseradish peroxidase (HRP) (Santa Cruz Biotechnology, sc-2357), or goat anti-mouse HRP (Santa Cruz Biotechnology, sc-2005), all diluted 1:1000-2500 in TBS-T plus 5% dry milk. Secondary antibodies were detected with ECL Western Blotting Detection Reagents (GE, Healthcare, RPN2106) for 1 min at room temperature, and developed using Biomax Light Film (Carestream, 178-8207).

### Immunofluorescence microscopy

*R. parkeri* infections were carried out in 24-well plates with sterile circle 12-mm coverslips (Thermo Fisher Scientific, 12-545-80). To initiate infection, 30%-purified bacteria were diluted in cell culture media at room temperature to a MOI of 0.01 for Vero cells, and a MOI of 0.1 for BMDMs. Bacteria were centrifuged onto cells at 300*g* for 5 min at room temperature and subsequently incubated at 33 °C. Next, infected cells were fixed for 10 min at room temperature in pre-warmed (37 °C) 4% PFA diluted in PBS, pH 7.4, then washed 3 times with PBS. Primary antibodies were the following: for staining with the guinea pig polyclonal anti-p62 antibody (Fitzgerald, 20R-PP001; 1:500 dilution), mouse polyclonal anti-NDP52 antibody (Novus Biologicals, H00010241-B01P; 1:100 dilution), a rabbit anti-*Rickettsia* I7205 antibody (1:500 dilution; gift from Ted Hackstadt), or anti-polyubiquitin FK1 antibody (1:250 dilution), cells were permeabilized with 0.5% Triton-X100 for 5 min prior to staining. For staining with mouse monoclonal anti-OmpA 13-3 antibody (1:5000 dilution), anti-OmpB antibody (*11*) (1:1000 dilution), or rabbit anti-O-antigen serum (*15*) (1:500 dilution), infected cells were post-fixed in methanol for 5 min at RT (no Triton-X). Cells were then washed 3 times with PBS and incubated with the primary antibodies for 30 min at RT. To detect the primary antibodies, secondary goat anti-rabbit Alexa-568 (Invitrogen, A11036), goat anti-mouse Alexa-568 (Invitrogen, A11004), or goat anti-guinea pig Alexa-568 (Invitrogen, A11075), Alexa 488 anti-rabbit antibody (Invitrogen, A11008; 1:500 dilution), Alexa 488 anti-mouse antibody (Invitrogen, A11001) antibodies were incubated at room temperature for 30 min (all 1:500 in PBS with 2% BSA). Images were captured as 15-25 z-stacks (0.1-μm step size) on a Nikon Ti Eclipse microscope with a Yokogawa CSU-XI spinning disc confocal with 100x (1.4 NA) Plan Apo objectives, and a Clara Interline CCD Camera (Andor Technology) using MetaMorph software (Molecular Devices). Images were processed using ImageJ using z-stack average maximum intensity projections and assembled in Adobe Photoshop. For quantification of the percentage of bacteria with pUb and p62, only bacteria that co-localized with rim-like patterns of the respective marker were scored as positive for staining. To quantify pUb, p62, NDP52, OmpB, and OmpA signal per bacteria, z-stacks were projected as stated above, and the edges of individual bacteria were marked by the freehand region of interest (ROI) function in ImageJ. Subsequently, the average pixel intensity within that ROI was measured. pUb/p62/NDP52 signal intensities were calculated by subtracting the average pUb/p62/NDP52 signal of WT bacteria from the pUb/p62/NDP52-value of each bacterium. OmpB signal intensity was calculated by subtracting the average OmpB-signal of *ompB*^*STOP*^::tn bacteria from the OmpB-value of each bacterium. OmpA signal intensity was calculated by subtracting the average background-signal (areas with no bacteria) from the OmpA-value of each bacterium.

### Sample preparation for mass spectrometry to determine the lysine methylome

5×10^7^ gradient-purified WT (Passage 6), *pkmt1*::tn (P4) and *pkmt2*::tn (P4) bacteria were centrifuged at 11,000*g* for 3 min. Each pellet was resuspended in 50 μL Tris (10 mM)-EDTA (10 mM), pH 7.6, and incubated for 45 min in a 45 °C water bath. Bacterial surface fractions were recovered from the supernatant after centrifugation at 11,000*g* for 3 min. Pellet was resuspended as above and incubated for additional 45 min at 45 °C before resuspension in 50 μL Tris (10 mM)-EDTA (10 mM). Both pellet and surface fractions were boiled at 95 °C for 10 min. Samples were cooled to RT prior to addition of 20 μL 50 mM NH_4_HCO_3_, pH 7.5, and 50 μL of a 0.2% solution of RapiGest (diluted in NH_4_HCO_3,_ Waters, 186001861). Next, samples were heated at 80 °C for 15 min and cooled to RT before addition of 1 μg of trypsin (Promega, V511A). Samples were digested at 37 °C overnight. To hydrolyze the RapiGest, 20 μL of 5% trifluoroacetic acid (TFA) was added to samples which were incubated at 37 °C for 90 min prior centrifugation at 15000*g* for 25 min at 4 °C. Samples were desalted using C18 OMIX tips (Agilent Technologies, A57003100) according to the manufacturer’s instructions and sample volume was decreased to 20 μL using a SpeedVac vacuum concentrator. Samples were stored at 4 °C prior to analysis.

### TUBE assay and sample preparation for mass spectrometry

To enrich for polyubiquitylated proteins, 3×10^8^ PFUs of “purified” WT (P6), *ompB*^*STOP*^::tn (P6) and *pkmt1*::tn (P4) and *pkmt2*::tn (P3) bacteria were centrifuged at 14,000*g* for 3 min at room temperature. Next, to release the surface protein fraction, the bacterial pellets were resuspended in lysis buffer (50 mM Tris-HCI, pH 7.5, 150 mM NaCl, 1 mM EDTA and 10% glycerol), supplemented with 0.0031% v/v lysonase (Millipore, 71230), the deubiquitylase inhibitor PR619 (Life Sensor, SI9619) at a final concentration of 20 μM, and 0.8% w/v octyl β-D-glucopyranoside (Sigma, O8001), and incubated on ice for 10 min with occasional pipetting of samples to break pellet into smaller pieces. Subsequently, the lysate was cleared by centrifugation at 14,000*g* at 4 °C for 5 min and incubated with equilibrated agarose TUBE-1 (Life Sensor, UM401) for 3 h, at 4 °C. After binding of polyubiquitylated proteins to TUBE-1, agarose beads were washed 1 time with TBS supplemented with 0.05% Tween and 5 mM EDTA, and subsequently 3 times with TBS only (no Tween or EDTA) and centrifuged at 5,000*g* for 5 min. To prepare samples for MS analysis, enriched proteins were digested at 37 °C overnight on agarose beads in RapiGest SF solution (Waters, 186001861) supplemented with 0.75 μg trypsin (Promega, V511A). The reaction was stopped using 1% TFA (Sigma, T6508). Octyl β-D-glucopyranoside was extracted using water-saturated ethyl acetate (Sigma, 34858). Prior to submission of samples for mass spectrometry analysis, samples were desalted using C18 OMIX tips (Agilent Technologies, A57003100), according to the manufacturer’s instructions.

### Liquid chromatography-mass spectrometry

Samples of proteolytically digested proteins were analyzed using a Synapt G2-S*i* ion mobility mass spectrometer that was equipped with a nanoelectrospray ionization source (Waters). The mass spectrometer was connected in line with an Acquity M-class ultra-performance liquid chromatography system that was equipped with trapping (Symmetry C18, inner diameter: 180 μm, length: 20 mm, particle size: 5 μm) and analytical (HSS T3, inner diameter: 75 μm, length: 250 mm, particle size: 1.8 μm) columns (Waters). Data-independent, ion mobility-enabled, high-definition mass spectra and tandem mass spectra were acquired in the positive ion mode (*23-25*). Raw data acquisition was controlled using MassLynx software (version 4.1), and tryptic peptide identification and relative quantification using a label-free approach (*26, 27*) were performed using Progenesis QI for Proteomics software (version 4.0, Waters). Raw data were searched against *Rickettsia parkeri* and *Chlorocebus sabaeus* protein databases (National Center for Biotechnology Information, NCBI) to identify tryptic peptides, allowing for up to three missed proteolytic cleavages, with diglycine-modified lysine (i.e., ubiquitylation remnant) and methylated lysine as variable post-translational modifications. Calculation of the percentage of lysine methylation (mono, di, tri or unmethylated), for each bacterial strain, was performed by dividing the abundance of a residue/protein bearing a modification by the total abundance and multiplying by 100. Data-dependent analysis was performed using an UltiMate3000 RSLCnano liquid chromatography system that was connected in line with an LTQ-Orbitrap-XL mass spectrometer equipped with a nanoelectrospray ionization source, and Xcalibur (version 2.0.7) and Proteome Discoverer (version 1.3, Thermo Fisher Scientific, Waltham, MA) software, as described elsewhere (*28*).

### Localization of tagged ubiquitin and ubiquitin pull-downs

To assess localization of 6xHis-ubiquitin during infection, confluent Vero cells grown in 24-well plates with coverslips were transfected with 2 μg of pCS2-6xHis-ubiquitin plasmid DNA using Lipofectamine 2000 (Invitrogen, 11668-019) for 6 h in Opti-MEM (Gibco, 31985-070). Subsequently, media were exchanged to media without transfection reagent, and cells were incubated overnight at 37 °C and 5% CO_2_. The following day (∼16 h after transfection), transfected cells were infected with purified WT or mutant bacteria at a MOI of 1. At 28 h.p.i., infected cells were fixed with 4% PFA diluted in PBS, pH 7.4 for 10 min, then washed 3 times with PBS. Primary anti-6xHis monoclonal mouse antibody (Clontech, 631212, diluted 1:1,000) was used to detect 6xHis-ubiquitin in samples permeabilized with 0.5% Triton-X100, and a goat anti-mouse Alexa-568 (Invitrogen, A11004) to detect the primary 6xHis antibody. Hoechst (Thermo Scientific, 62249, diluted 1:2500) was used to detect host and bacterial DNA. Samples were imaged as already described.

For ubiquitin pull-downs, confluent Vero cells grown in 6-well plates were transfected and infected as described above. At 28 h.p.i., cells were washed once with 1x PBS, pH 7.4, and subsequently lysed in urea lysis buffer (8 M urea, 50 mM Tris-HCI, pH 8.0, 300 mM NaCI, 50 mM Na_2_HPO_4_, 0.5% Igepal CA-630 (Sigma, I8896)) for 20 min at RT. Subsequently, samples were sonicated, and lysate was cleared by centrifugation at 15000*g* for 15 min at room temperature. Prior to incubation with Ni-NTA resin, an aliquot was saved for the input sample. 6xHis-ubiquitin conjugates were purified by incubation and rotation with Ni-NTA resin for 3 h, at RT, in the presence of 10 mM imidazole. Beads were washed 3 times with urea lysis buffer and 1 time with urea lysis buffer lacking Igepal CA-630. Ubiquitin conjugates were eluted at 65 °C for 15 min in 2x Laemmli buffer containing 200 mM imidazole and 5% 2-mercaptoetanol (Sigma, M6250), vortexed for 90 seconds, and centrifuged at 5000*g* for 5 min at RT. Eluted and input proteins were detected by SDS-PAGE followed by western blotting, as described above.

### Animal experiments

Animal research using mice was conducted under a protocol approved by the UC Berkeley Institutional Animal Care and Use Committee (IACUC) in compliance with the Animal Welfare Act. The UC Berkeley IACUC is fully accredited by the Association for the Assessment and Accreditation of Laboratory Animal Care International and adheres to the principles of the Guide for the Care and Use of Laboratory Animals. Infections were performed in a biosafety level 2 facility. Mice were age-matched between 8 and 18 weeks old. Mice were selected for experiments based on their availability, regardless of sex. All mice were healthy at the time of infection and were housed in microisolator cages and provided chow and water. Littermates of the same sex were randomly assigned to experimental groups. For infections, *R. parkeri* was prepared by diluting 30%-prep bacteria into cold sterile 1x PBS to 5×10^6^ PFU per 200 mL. Bacterial suspensions were kept on ice during injections. Mice were exposed to a heat lamp while in their cages for approximately 5 min, and then each mouse was moved to a mouse restrainer (Braintree, TB-150 STD). The tail was sterilized with 70% ethanol, and 200 µL of bacterial suspensions were injected using 30.5-gauge needles into the lateral tail vein. Body temperatures were monitored using a rodent rectal thermometer (BrainTree Scientific, RET-3). Mice were monitored daily for clinical signs of disease, such as hunched posture, lethargy, or scruffed fur. If a mouse displayed severe signs of infection, as defined by a reduction in body temperature below 90 °F or an inability to move around the cage normally, the animal was immediately and humanely euthanized using CO_2_ followed by cervical dislocation, according to IACUC-approved procedures (*16*).

### Statistical analysis, experimental variability and reproducibility

Statistical parameters and significance are reported in the legends. Data were considered to be statistically significant when p < 0.05, as determined by a one-way ANOVA with Dunnett’s post-hoc test, a Kruskal-Wallis test with Dunn’s post-hoc test, a Brown-Forsyth and Welch ANOVA with Dunnett’s post-hoc test, or a two-way ANOVA (all two-sided). Statistical analysis was performed using PRISM 6 software (GraphPad Software). If not otherwise described, *n* indicates the number of independent biological experiments executed at different times.

## References and Notes

1. F. Randow, R. J. Youle, Self and nonself: how autophagy targets mitochondria and bacteria. Cell Host Microbe 15, 403–411 (2014).

2. A. J. Perrin, X. Jiang, C. L. Birmingham, N. S. So, J. H. Brumell, Recognition of bacteria in the cytosol of Mammalian cells by the ubiquitin system. Curr Biol 14, 806–811 (2004).

3. S. J. L. van Wijk et al., Linear ubiquitination of cytosolic Salmonella Typhimurium activates NF-kappaB and restricts bacterial proliferation. Nat Microbiol 2, 17066 (2017).

4. J. Noad et al., LUBAC-synthesized linear ubiquitin chains restrict cytosol-invading bacteria by activating autophagy and NF-kappaB. Nat Microbiol 2, 17063 (2017).

5. E. D. Case et al., The Francisella O-antigen mediates survival in the macrophage cytosol via autophagy avoidance. Cell Microbiol 16, 862–877 (2014).

6. J. Huang, J. H. Brumell, Bacteria-autophagy interplay: a battle for survival. Nat Rev Microbiol 12, 101-114 (2014).

7. Y. T. Zheng et al., The adaptor protein p62/SQSTM1 targets invading bacteria to the autophagy pathway. J Immunol 183, 5909–5916 (2009).

8. T. L. Thurston, G. Ryzhakov, S. Bloor, N. von Muhlinen, F. Randow, The TBK1 adaptor and autophagy receptor NDP52 restricts the proliferation of ubiquitin-coated bacteria. Nat Immunol 10, 1215–1221 (2009).

9. E. Fiskin, T. Bionda, I. Dikic, C. Behrends, Global analysis of host and bacterial ubiquitinome in response to Salmonella Typhimurium infection. Mol Cell 62, 967–981 (2016).

10. M. Polajnar, M. S. Dietz, M. Heilemann, C. Behrends, Expanding the host cell ubiquitylation machinery targeting cytosolic Salmonella. EMBO Rep 18, 1572–1585 (2017).

11. P. Engström et al., Evasion of autophagy mediated by Rickettsia surface protein OmpB is critical for virulence. Nat Microbiol 4, 2538–2551 (2019).

12. S. Lanouette, V. Mongeon, D. Figeys, J. F. Couture, The functional diversity of protein lysine methylation. Mol Syst Biol 10, 724 (2014).

13. R. L. Lamason, N. M. Kafai, M. D. Welch, A streamlined method for transposon mutagenesis of Rickettsia parkeri yields numerous mutations that impact infection. PLoS One 13, e0197012 (2018).

14. B. G. Spratt, A. B. Pardee, Penicillin-binding proteins and cell shape in E. coli. Nature 254, 516–517 (1975).

15. H. K. Kim, R. Premaratna, D. M. Missiakas, O. Schneewind, Rickettsia conorii O antigen is the target of bactericidal Weil-Felix antibodies. Proc Natl Acad Sci U S A 116, 19659–19664 (2019).

16. T. P. Burke et al., Inflammasome-mediated antagonism of type I interferon enhances Rickettsia pathogenesis. Nat Microbiol 5, 688–696 (2020).

17. D. C. H. Yang, A. H. Abeykoon, B. E. Choi, W. M. Ching, P. B. Chock, Outer membrane protein OmpB methylation may mediate bacterial virulence. Trends Biochem Sci 42, 936–945 (2017).

18. A. H. Abeykoon et al., Structural insights into substrate recognition and catalysis in outer membrane protein B (OmpB) by protein-lysine methyltransferases from Rickettsia. J Biol Chem 291, 19962–19974 (2016).

19. A. Abeykoon et al., Multimethylation of Rickettsia OmpB catalyzed by lysine methyltransferases. J Biol Chem 289, 7691–7701 (2014).

20. A. H. Abeykoon et al., Two protein lysine methyltransferases methylate outer membrane protein B from Rickettsia. J Bacteriol 194, 6410–6418 (2012).

21. N. F. Noriea, T. R. Clark, T. Hackstadt, Targeted knockout of the Rickettsia rickettsii OmpA surface antigen does not diminish virulence in a mammalian model system. mBio 6, (2015).

22. R. L. Lamason et al., Rickettsia Sca4 Reduces Vinculin-Mediated Intercellular Tension to Promote Spread. Cell 167, 670–683 e610 (2016).

23. U. Distler et al., Drift time-specific collision energies enable deep-coverage data-independent acquisition proteomics. Nat Methods 11, 167–170 (2014).

24. R. S. Plumb et al., UPLC/MS(E); a new approach for generating molecular fragment information for biomarker structure elucidation. Rapid Commun Mass Spectrom 20, 1989–1994 (2006).

25. P. V. Shliaha, N. J. Bond, L. Gatto, K. S. Lilley, Effects of traveling wave ion mobility separation on data independent acquisition in proteomics studies. J Proteome Res 12, 2323–2339 (2013).

26. S. Nahnsen, C. Bielow, K. Reinert, O. Kohlbacher, Tools for label-free peptide quantification. Mol Cell Proteomics 12, 549–556 (2013).

27. K. A. Neilson et al., Less label, more free: approaches in label-free quantitative mass spectrometry. Proteomics 11, 535–553 (2011).

28. C. I. Carlströmet al., (Per)chlorate-reducing bacteria can utilize aerobic and anaerobic pathways of aromatic degradation with (per)chlorate as an electron acceptor. mBio 6, (2015).

29. R. L. Lamason et al., Rickettsia Sca4 Reduces Vinculin-Mediated Intercellular Tension to Promote Spread. Cell 167, 670–683 e610 (2016).

30. U. Distler et al., Drift time-specific collision energies enable deep-coverage data-independent acquisition proteomics. Nat Methods 11, 167–170 (2014).

31. R. S. Plumb et al., UPLC/MS(E); a new approach for generating molecular fragment information for biomarker structure elucidation. Rapid Commun Mass Spectrom 20, 1989–1994 (2006).

32. P. V. Shliaha, N. J. Bond, L. Gatto, K. S. Lilley, Effects of traveling wave ion mobility separation on data independent acquisition in proteomics studies. J Proteome Res 12, 2323–2339 (2013).

33. S. Nahnsen, C. Bielow, K. Reinert, O. Kohlbacher, Tools for label-free peptide quantification. Mol Cell Proteomics 12, 549–556 (2013).

34. K. A. Neilson et al., Less label, more free: approaches in label-free quantitative mass spectrometry. Proteomics 11, 535–553 (2011).

35. C. I. Carlström et al., (Per)chlorate-reducing bacteria can utilize aerobic and anaerobic pathways of aromatic degradation with (per)chlorate as an electron acceptor. mBio 6, e02287–14 (2015).

